# Conserved multiheme cytochrome machinery for extracellular electron transfer is widespread and transcriptionally active across deep peat profiles

**DOI:** 10.64898/2026.05.27.727921

**Authors:** Giulia Fiorito, Taylor Priest, Bledina Dede, Hanna Zehnle, Martin H. Schroth, Michael Sander, Marie C. Schoelmerich

## Abstract

Northern peatlands store approximately one-third of global soil organic carbon, yet the anaerobic respiratory pathways governing carbon turnover remain unclear. In ombrotrophic bogs, the scarcity of inorganic terminal electron acceptors (TEAs) and high CO_2_:CH_4_ ratios indicate that methanogenesis alone cannot account for the observed CO_2_ production. Peat particulate organic matter (POM) has been proposed as an alternative TEA, but whether resident microorganisms encode and express extracellular electron transfer (EET) machinery required to use such extracellular TEAs remains unknown. Using depth-resolved metagenomics and metatranscriptomics across peat profiles from four ombrotrophic Swedish bogs, we identified conserved EET machinery in dominant yet uncultured Acidobacteriota and Verrucomicrobiota, comprising multiheme cytochromes and outer-membrane porins arranged in syntenic gene clusters. This machinery was transcriptionally active up to several meters’ depth, alongside broader anaerobic respiratory pathways, while methane-cycling processes were more prominent in the upper layers. These results provide systematic genomic and transcriptomic evidence for EET capacity in peatland microorganisms, establishing a molecular foundation for EET-based respiration and its potential role in suppressing methane formation and emissions.

## Introduction

Northern peatlands represent one of Earth’s largest terrestrial carbon reservoirs. In these systems, long-term carbon accumulation occurs because photosynthetic CO_2_ fixation by peat-forming vegetation proceeds faster than microbial organic matter decomposition under the waterlogged and acidic conditions that characterize these ecosystems. At the same time, they are globally significant sources of methane (CH_4_), contributing an estimated ∼5-11% of annual atmospheric emissions^1^. Ombrotrophic bogs, a major subtype of northern peatlands, are fed exclusively by precipitation, making them particularly nutrient-poor. Oxygen is restricted to the acrotelm, a thin aerobic surface layer with active vegetation, while the underlying catotelm is permanently waterlogged and anoxic. Inorganic terminal electron acceptors (TEAs) such as nitrate, sulfate, iron and manganese are scarce, suggesting methanogenesis as the dominant respiratory process^2^. This expectation is consistent with the widespread occurrence of bog-adapted methanogens, such as the archaeon *Candidatus* Methanoflorens stordalenmirensis (Bog-38) in northern peatlands^3,4^. Complete methanogenic degradation of organic matter with a net oxidation state of carbon close to zero is expected to yield CO_2_ and CH_4_ in an approximately 1:1 molar ratio. Yet, dissolved CO_2_:CH_4_ concentration ratios in porewater consistently exceed unity, often by orders of magnitude, even at sites where concentrations of canonical inorganic TEAs are vanishingly low^3,5,6^. Excess CO_2_ formation therefore suggests anaerobic respiration in which reducing equivalents are diverted to an as-yet unidentified TEA, thereby competitively suppressing methanogenesis and thus lowering CH_4_ emissions.

Recent studies have begun to identify seemingly overlooked TEAs. Metagenomic surveys have revealed that Acidobacteriota, which dominate northern bog communities and encode extensive enzymatic machinery for plant organic matter degradation^7^, carry choline-sulfatase (*betC*), which may liberate sulfate from *Sphagnum*-derived choline-O-sulfate^3^. This intracellular enzyme would generate sulfate as TEA supply directly from organic precursors. Separately, electrochemical studies have shown that peat particulate organic matter (POM) carries redox-active chemical moieties, tentatively quinones, that reversibly accept electrons under anoxic conditions and are reoxidized upon transient oxygen inputs, such as water table fluctuations and oxygen transport through plant aerenchyma^8,9^. POM thus represents a plausible extracellular and regenerable electron acceptor, but its sustained function as a TEA requires continuous reoxidation. Together, these observations suggest that peat organic matter may underpin electron acceptor availability in ombrotrophic bogs through two distinct routes: indirectly, as a precursor to inorganic and soluble TEAs, and directly, as redox-active extracellular electron acceptor. Reduction of insoluble or extracellular TEAs such as POM would require electrons to be transferred across the cell envelope in a process termed extracellular electron transfer (EET).

The molecular machinery mediating such EET has been extensively characterized in representatives of the genera *Shewanella* and *Geobacter*. In these organisms, multiheme cytochromes (MHCs) mediate electron transfer to extracellular and insoluble terminal acceptors, including Fe(III), Mn(IV) and humic substances^10,11^. In other systems, such as marine sediments, cable bacteria of the candidate genera *Electrothrix* and *Electronema* perform centimeter-scale electron transfer through filamentous structures, coupling spatially separated oxidation and reduction half-reactions across steep redox gradients^12^. These examples illustrate the diversity of microbial strategies for transferring electrons beyond the cell. Whether such electron transfer machinery exists in the microorganisms dominant in ombrotrophic bogs, however, remains unclear. Prior genomic work has noted porin-MHC gene clusters in Acidobacteriota^13^ and hinted at EET potential in humic-rich freshwater systems^14^, but no systematic characterization of EET machinery has been reported in peatland microbiomes. Thus, the microorganisms capable of EET in these ecosystems, and the molecular machinery enabling this process, remain unknown.

Here, we integrate field sampling and biogeochemical measurements with 16S rRNA gene amplicon, metagenomic and metatranscriptomic sequencing across four Swedish bogs to address three questions: 1. which microorganisms dominate across peat profiles, including the deepest cores extending up to 7 m, and how does community structure vary from the acrotelm to the deep catotelm? 2. do dominant lineages encode molecular machinery consistent with EET, and what are the structural and phylogenetic characteristics of these systems? 3. is this machinery transcriptionally active throughout the peat profile, including at depths where inorganic TEAs are undetectable? We show that the Halobacteriota archaeon Bog-38 (*Candidatus* Methanoflorens stordalenmirensis) dominates the methanogen community yet co-occurs with active methane oxidizers even in deep layers. Acidobacteriota and Verrucomicrobiota encode exceptionally large and diverse MHC repertoires and harbor four conserved EET gene cluster architectures, each comprising MHCs and outer-membrane porins, whose predicted structures are reminiscent of characterized EET conduits in *Shewanella* and *Geobacter,* yet entirely divergent in sequence. All four systems are actively transcribed *in situ* across all sampled depths, providing the first systematic evidence for convergently evolved EET machinery in dominant, uncultured peat bacteria. These findings suggest that EET may represent a widespread but previously unrecognized respiratory strategy in ombrotrophic peatlands, with redox-active POM as a plausible candidate TEA.

## Results

### Depth-structured peat microbial communities suggest alternative electron-flow pathways

To assess microbial community structure and biogeochemical cycling in northern peatlands, we sampled vertical peat profiles in four pristine ombrotrophic bogs in Värmland, central Sweden: Björsmossen (BM), Havsjömossen (HV), Lungsmossen (LM) and Norra Romyren (NR) (**ED Figure 1A**). These *Sphagnum*-dominated raised bogs developed in the Holocene approximately 6,000 years BP^15^ and are characterized by persistent anoxic conditions in the catotelm that shape steep redox gradients and structure microbial communities. Cores spanning the upper 0-60 cm of the peat were collected from three spatial replicates in BM, NR and LM each to capture near-surface variation (spatial profiles), while deep cores from all four peatlands were used to elucidate depth-related patterns (deep profiles). Porewater analyses of the spatial profiles confirmed highly acidic (pH 3.8 - 4.2), nutrient-poor conditions with low conductivity (30-60 µS cm^−1^) (**ED Figure 1B**,^8,9^). Low organic acid concentrations (propionate ∼0.7 mM, acetate ∼20 μM; **ED Figure 1B**) indicated rapid consumption of fermentation products. Inorganic TEAs, such as nitrate (NO_3_^−^), sulfate (SO_4_^2−^), total manganese (Mn) and total iron (Fe), were scarce or below detection limits (**Figure 1A**). Notably, under anoxic porewater conditions measured, dissolved Fe and Mn are expected to be predominantly Fe(II) and Mn(II), implying limited Fe(III)/Mn(IV) availability as TEAs. Despite this canonical inorganic TEA limitation, dissolved CO_2_:CH_4_ molar concentration ratios ranged from 2.0-16.5 across sites and depths, indicating an alternative anaerobic respiratory pathway beyond methanogenesis (**Figure 1A**).

**Figure 1.**
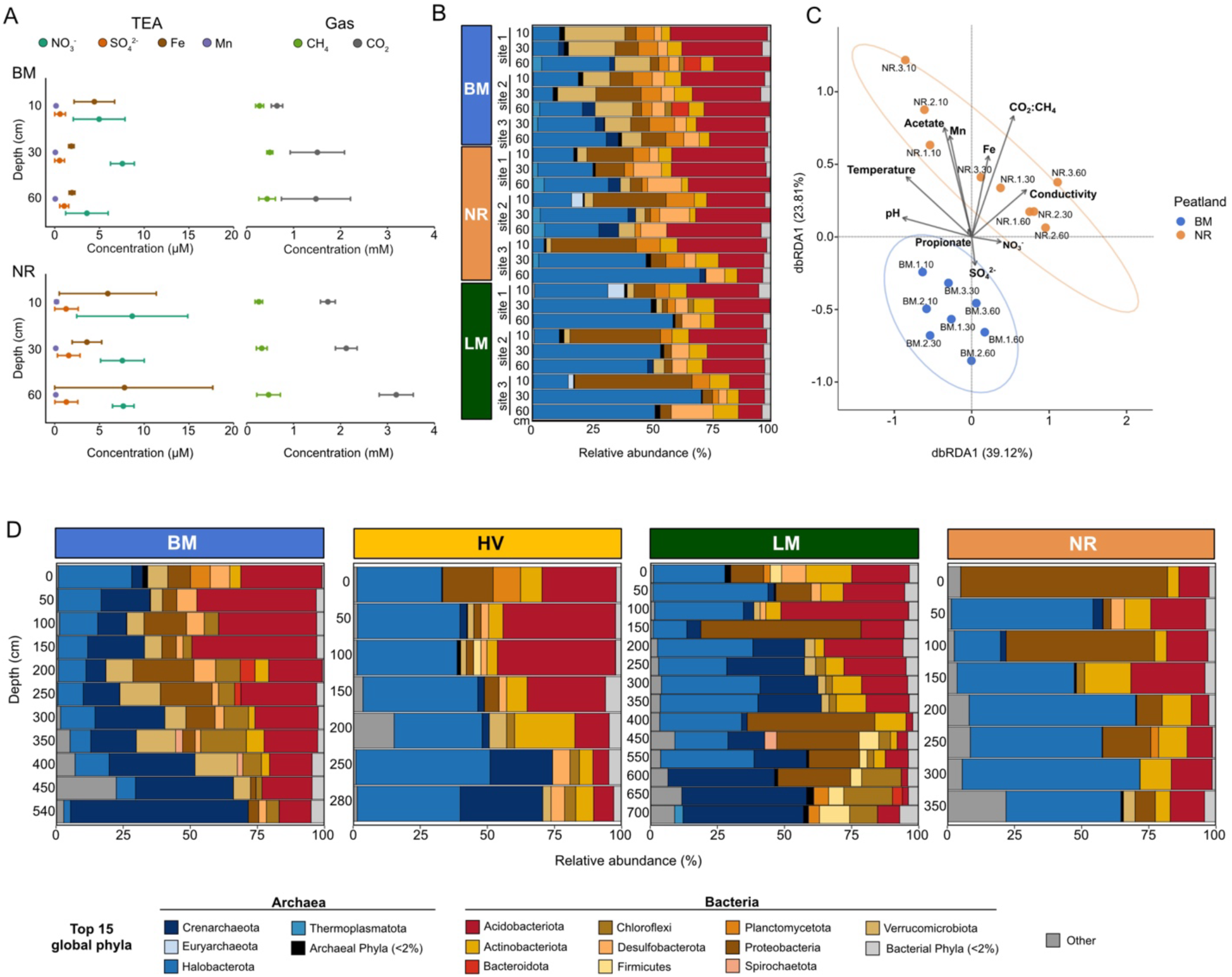
Microbial communities and biogeochemistry along peat depth profiles. **A**. Depth profiles showing concentrations of the terminal electron acceptors (TEAs) (NO_3_^−^, SO_4_^2−^, Mn, Fe) and gases (CH_4_,CO_2_) for BM and NR in spatial profiling, shown as the mean values across sites 1-3 for each depth (10, 30, 60cm). Bars indicate standard deviation. **B**. Bar plot showing the relative abundance (%) of the top 15 global phyla (spatial and deep sampling) shown for spatial samples at different soil depths (10, 30, 60 cm) and three sites in BM (blue), NR (orange), LM (green). Phylum-level legend is shown in **D**. **C**. Distance-based redundancy analysis (dbRDA) showing relationship between microbial community profiles with 16S rRNA gene amplicon sequencing and biogeochemical measurements for BM and NR in spatial profiling. Arrows indicate environmental variables (pH, temperature, Conductivity, CO_2_:CH_4_, acetate, propionate, total manganese (Mn), total iron (Fe), sulfate (SO_4_^2−^) and nitrate (NO_3_^−^)). Blue (BM) and orange (NR) points indicate peat microbial samples, while in the label above the points, 1,2 and 3 indicate sites and 10, 30 and 60 cm the corresponding depth per sample, respectively. **D**. Relative abundance (%) of the top 15 global phyla from spatial and deep sampling in the deep profiles across four peatlands: BM (blue), HV (yellow), LM (green), NR (orange). Color legend includes both bacterial and archaeal phyla, including taxa representing <2% of the total relative abundance. Other: other phyla.

16S rRNA gene amplicon sequencing showed that microbial communities varied strongly with depth. In the spatial profiles, richness, evenness and Shannon diversity generally declined towards 60 cm, along with increasing similarity in community composition at consecutive depths, as evidenced by changes in Bray-Curtis and Jaccard distances (**ED Figure 1C**). However, deep profiles revealed additional, site-specific shifts in diversity and composition in deeper layers. Notably, microbial diversity increased and community composition shifted markedly below 450 cm depth at the LM site and between 250 - 350 cm at NR site (**ED Figure 1C)**. These patterns identify depth as a major axis structuring peat microbial communities but also highlight that the magnitude and direction of such changes is heterogeneous across peatlands.

To determine how microbial community composition varied in relation to biogeochemical conditions, we focused on the upper peat layers for which porewater chemistry was measured. Microbial communities in the spatial profiles were dominated by both bacterial and archaeal lineages characteristic of acidic, organic-rich peat. Acidobacteriota (26.9%±9.11%) and Halobacterota (Halobacteriota in GTDB^16^ database: 30.5%±19.1%) comprised the most abundant fractions, with Proteobacteria (Pseudomonadota in GTDB database: 11.4%±12.6%), Verrucomicrobiota (6.34%±6.68%) and Desulfobacterota (6.03%±4.26%), alongside the archaeal phylum Crenarchaeota (Thermoproteota in GTDB database: 2.05%±1.93%) which was also consistently detected (**Figure 1B**). To assess how compositional shifts across the upper 60 cm related to measured biogeochemical gradients, we performed distance-based redundancy analysis (dbRDA). This revealed a distinct separation between samples from BM and those from NR and LM when considering the environmental variables measured across all three peatlands (pH, temperature, conductivity, CO_2_:CH_4_ ratios). Within NR and LM, conductivity and gas concentration ratios were more strongly associated with samples below 30 cm, while temperature was more closely associated with the upper 10 cm samples. By contrast, BM samples were primarily associated with pH (**Figure 1C; ED Figure 1D**). Restricting the analysis to BM and NR, where the full porewater geochemical dataset of measured organic acids and potential canonical inorganic TEAs was available, further identified Fe, Mn and acetate as additional contributors to the separation, whereas propionate, SO_4_^2−^and NO_3_^−^ showed weaker associations (**Figure 1C**).

Deep peat profiles revealed that the dominant microbial lineages detected in the upper layers persisted across the peat profile, but with marked changes in their relative abundance at depth (**Figure 1D**). Of particular note, Archaea represented a larger fraction of communities in deep profiles than in spatial profiles (42.1%±16.1% versus 34.6%±18.7%). Across the peat profiles, the most abundant archaeal lineage was the Halobacterota clade Rice Cluster II (also referred to as Bog-38 or *Ca.* Methanoflorens), a predicted methanogen that is prevalent in other northern peatland locations, such as SPRUCE^3^ and Stordalen Mire^4^. However, in the deepest layers, canonical methanogens of the Halobacterota represented an increasingly small fraction of the community (∼1%), while members of the Crenarchaeota became dominant, suggesting substrate limitation for methanogenesis^17^ and a shift towards alternative anaerobic pathways. Underpinning this transition were uncultivated members of Bathyarchaeia, which became particularly prominent below 5 m depth at both BM and LM sites. Thus, while many major surface-associated lineages persisted across depth, community structure shifted markedly in the deepest peat layers, largely reflecting a transition from Halobacterota-rich to Crenarchaeota-dominated archaeal assemblages, consistent with increasing peat age and organic matter recalcitrance at depth^15^.

Together, these results identify depth as a major factor structuring microbial communities across ombrotrophic bog peat profiles. In the upper 60 cm, this structure emerged under acidic, nutrient-poor and canonical inorganic TEA-limited conditions, yet dissolved CO_2_:CH_4_ concentration ratios consistently exceeded unity, indicating that reducing equivalents were diverted to TEAs beyond the measured inorganic canonical TEA pool. Deep peat profiles extended this depth-structured pattern across the full peat column, with the deepest layers marked by a shift towards Crenarchaeota-dominated archaeal assemblages.

### Metagenomic profiling captures peat community structure and enables functional analysis across depth layers

To link microbial community composition to metabolic function across depth, we performed shotgun metagenomic and metatranscriptomic sequencing on a subset of 32 samples from both shallow and deep layers across the four bogs. Taxonomic profiling of metagenomic reads broadly recapitulated the community compositions inferred from 16S rRNA gene amplicons across sites and depths (**ED Figure 2A**), indicating that the metagenomic data captured the dominant lineages in the sampled communities. We next sought to contextualize the recovered communities with those from other well-studied northern peatlands, including SPRUCE (Minnesota, USA) and Stordalen Mire (Abisko, Sweden). Through whole-metagenome nucleotide composition comparisons, we observed a greater similarity between Värmland and SPRUCE bog communities than between Värmland and Stordalen Mire fen and palsa communities (**ED Figure 2B**). The latter presents minerotrophic rather than ombrotrophic conditions, which could be a reason for the observed compositional differences.

For a genome-resolved representation of the sampled peat communities, we generated a comprehensive metagenome-assembled genome (MAG) collection. Dereplication at species level yielded 1,081 medium- to high-quality MAGs (>50% completeness, <10% contamination), spanning 46 bacterial (910 MAGs) and 10 archaeal (171 MAGs) phyla. Of these, 472 MAGs met high-quality criteria (>90% completeness, <5% contamination), comprising 52 candidate novel species, which mainly belonged to the phylum Acidobacteriota. Although the MAG collection represented only 37.7% of metagenomic read pool, it represented all phyla exceeding 1% mean relative abundance in the 16S rRNA amplicon dataset; together these dominant phyla accounted for 94.5% of the amplicon-derived community abundance. Moreover, the recovered MAGs recruited ∼59% of corresponding metatranscriptomic reads, indicating that they captured a substantial fraction of the transcriptionally active community.

As the MAG catalogue captured a substantial but incomplete fraction of the community, we sought to incorporate information from unbinned metagenomic contigs to broaden functional insights across peat depths. For this, we combined genes from both MAGs and unbinned contigs to construct a non-redundant gene catalogue comprising 37 million gene clusters at 95% sequence identity. This catalogue was subsequently profiled across matched metagenomes and metatranscriptomes - using genome and gene copy number normalizations for cross-sample comparisons - enabling depth-resolved analyses on the abundance and transcriptional activity of genes, protein functions and metabolic pathways (**Figure 2; Supplementary discussion**).

**Figure 2.**
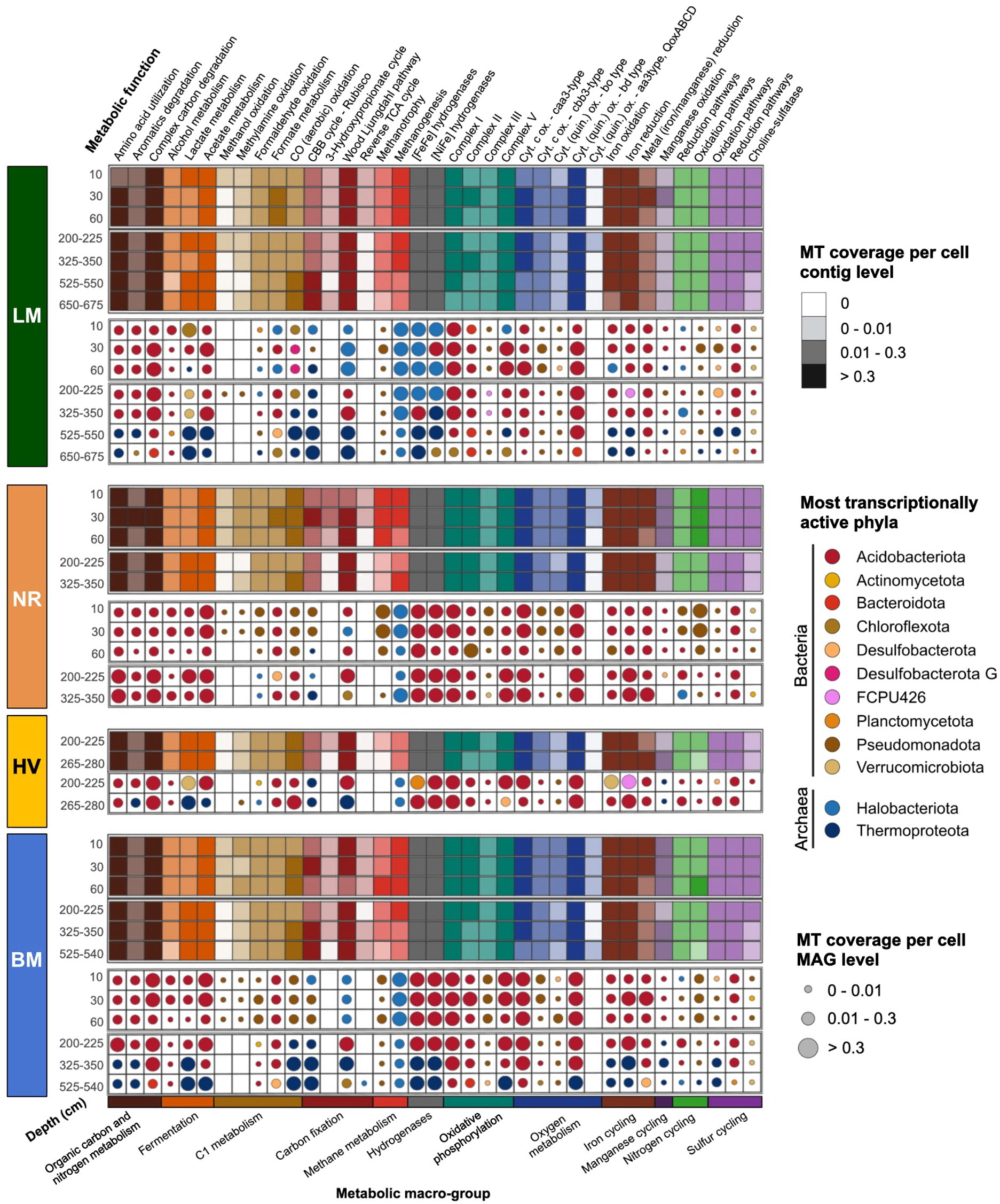
Depth-resolved expression of key metabolic function across four Swedish peatlands and the taxa contributing most to their expression. Integrated visualization of microbial expression functions across peatlands and depth layers (cm). For each peatland, the top panel (heatmap) shows functional expression at contig level as metatranscriptomic coverage normalized per genome equivalent. For each function, the expression of the corresponding marker genes was summed across samples and normalized by the number of markers per function. Colors indicate the metabolic macro-groups, while shading indicates MT coverage per cell levels (0, 0-0.01, 0.01-0.3, > 0.3). Bottom panel (bubble plot) indicates MAG-resolved taxonomic attribution of functional expression. For each combination of peatland, depth and function, function-level MT coverage per cell was first calculated per MAG (sum of marker expressions normalized by number of markers), then summed across MAGs within each phylum. Bubble color indicates the phylum with the highest MT coverage per cell per function, among the 15 most transcriptionally active phyla globally (archaea and bacteria in the legend), while bubble size encodes MT coverage per cell levels (0-0.01, 0.01-0.3, >0.3). Only phyla that emerged as the most expressing ones for the functions shown are displayed in the legend (12/15). Metabolic macro-group colors are shown below the x-axis. Absence of bubble indicates no detectable expression. LM: Lungsmossen; NR: Norra Romyren; HV: Havsjömossen; BM: Björsmossen.

### Depth-dependent activity patterns across key microbial metabolisms

Functional profiling revealed distinct depth-dependent patterns in the genetic potential and transcriptional activity of key microbial metabolisms. Across all depths, contig-level transcripts associated with organic carbon and nitrogen metabolism were consistently detected, indicating sustained heterotrophic activity throughout the peat profile. In upper layers, carbohydrate degradation was associated with CAZyme functions, primarily belonging to Acidobacteriota MAGs, alongside aromatic degradation genes (e.g., benzoyl-CoA reductase; *bcrABCD*). This observation is consistent with the breakdown of aromatic-rich peat organic matter. Sulfur cycling near the surface was marked by expression of choline-sulfatase (*betC*)^3^ by Verrucomicrobiota, particularly at NR 30 cm. Hydrogen metabolism in the same layers was dominated by Acidobacteriota expressing multiple hydrogenases of the [NiFe]- and [FeFe]-types (**ED Figure 3**).

With increasing depth, certain metabolic functions shifted: hydrogenase expression transitioned from Acidobacteriota in shallow layers to Thermoproteota at depth (**ED Figure 3**) indicating that H_2_ cycling (production via fermentation and oxidation) is a crucial process supporting anaerobic metabolisms across the peat profile. Wood-Ljungdahl pathway (WLP) gene (CO dehydrogenase/acetyl-CoA synthase complex, *cdhDE*; anaerobic carbon-monoxide dehydrogenase catalytic subunit, *cooS*) expression, although detected across all depths, was higher in the deeper layers, pointing to the coexistence of heterotrophic and autotrophic activity. In these deeper profiles, sulfur cycling genes for both oxidative and reductive pathways, were expressed by Acidobacteriota, Desulfobacterota and Thermoproteota. These genes included the dissimilatory sulfite reductase (*dsrAB*) and anaerobic sulfite reductase (*asrBC*). The latter was predominantly expressed by Verrucomicrobiota at LM 525-550 cm, Planctomycetota at LM 650-675 cm, and Thermoproteota and Halobacteriota at BM 525-550 cm. Together with widespread expression of iron-cycling genes, these patterns indicate that diverse anaerobic respiratory pathways remain active through deep layers where canonical inorganic TEAs are very scarce (**Figure 2; ED Figure 3**).

Given the central role of methane cycling in northern peatlands, we examined methane-cycling genes and their microbial mediators in greater detail. At the contig level, we observed higher transcriptional activity for both methanogenesis and methanotrophy in spatial compared to deep profiles (**Figure 2**). However, these signals varied among sites. The strongest methanogenic activity was observed at LM and the strongest methanotrophic activity was detected at NR. In deeper layers (>200 cm), the transcription of both processes generally decreased, with site-specific exceptions (e.g., NR 325-350 cm). Nevertheless, methane metabolism remained detectable across the communities (**Figure 2**). At the MAG level, we recovered 26 putative methanogen MAGs (harboring methyl-coenzyme M reductase, *mcrA*), and 8 potential methanotroph MAGs (carrying methane monooxygenase genes, *pmoA* and *mmoX*) (**ED Figure 4A,B,C; Supplementary discussion**). Methane oxidizing bacteria (MOB), mainly affiliated with Pseudomonadota, accounted for a mean relative abundance of ∼1.5% across all sites (**ED Figure 4A**). Notably, *Methylocystis-*affiliated *pmoA* transcripts were detected even in the deep layers, including BM 325-350 cm and LM 200-225 cm. This suggests that these populations may tolerate low-oxygen conditions at depth^18^ (**ED Figure 4B**). It also raises the possibility that methane oxidation, or another metabolism such as the oxidation of multicarbon compounds (e.g., acetate), persists under microoxic conditions in deep peat. In contrast, methanogens represented ∼14% of the total community. Bog-38 (4 MAGs) emerged as the most abundant and transcriptionally active putative methanogen across sites and depths (**ED Figure 4A,B**), co-occurring with active methanotrophs in both spatial (10-60 cm) and deep layers (>200 cm). The co-occurrence of active methanogens and methanotrophs across both spatial and deep profiles indicates that methane cycling persists throughout the peat profile, albeit with higher expression of the relevant genes in upper layers.

Together, these analyses reveal metabolically active microbial communities throughout the peat profile, spanning methane cycling in the upper layers to diverse anaerobic respiratory pathways at depth. However, persistently elevated dissolved CO_2_:CH_4_ concentration ratios and scarcity of measured inorganic TEAs point to additional anaerobic terminal respiratory processes in the deep catotelm and raise the question of what molecular machinery could mediate them.

### Multiheme cytochromes are widespread, diverse in sequence, and found in transcriptionally active MAGs

Having established that diverse anaerobic respiratory pathways are transcriptionally active across the peat profile, including in deep layers where canonical inorganic TEAs are scarce, we next investigated whether the dominant lineages encode molecular machinery consistent with EET. Given that EET to extracellular or insoluble acceptors is mediated by MHCs in characterized systems^11^, we set out to illuminate genomic evidence for the potential occurrence of such a process in our peat profiles.

Screening of all MAGs and unbinned contigs for proteins containing at least three heme-binding motifs, yielded 12,805 candidate MHCs. Across the candidates, heme content and density varied considerably, ranging from 3 to 116 motifs with a median of 1.13 motifs per 100 amino acids. This variability was underpinned by considerable sequence diversity, with candidate MHCs grouping into 1392 distinct, non-singleton clusters at 50% sequence identity. Among these, only 9 shared homology with MHCs from *Geobacter sulfurreducens* PCA^19^ and *Shewanella oneidensis* MR-1^20^. Thus, the diverse MHCs recovered from our peat profiles are largely distinct from experimentally characterized analogues.

To determine whether candidate MHCs may facilitate EET, we next predicted their cellular localization. A substantial fraction was predicted to span across the cell envelope, with ∼3% (379) predicted to be extracellular, ∼14% periplasmic and ∼17% associated with the cytoplasmic membrane, thus placing ∼34% of the repertoire in compartments relevant to electron transfer across or beyond the cell envelope.

Given these observations, we devised a credibility scoring framework leveraging heme density, predicted localization and functional annotation to prioritize candidates most likely to support EET. Through this, we identified 4,811 high-priority candidates (Tier 1; see **Methods**) exhibiting a high heme density (median 13 motifs; 3.7 motifs per 100 aa), a compact size (median 423 aa), an envelope-accessible localization and modular cytochrome architectures, which included *c*-type cytochrome, doubled CXXCH and quinol-oxidoreductase-associated domains (**Figure 3A**; **ED Figure 5**).

**Figure 3.**
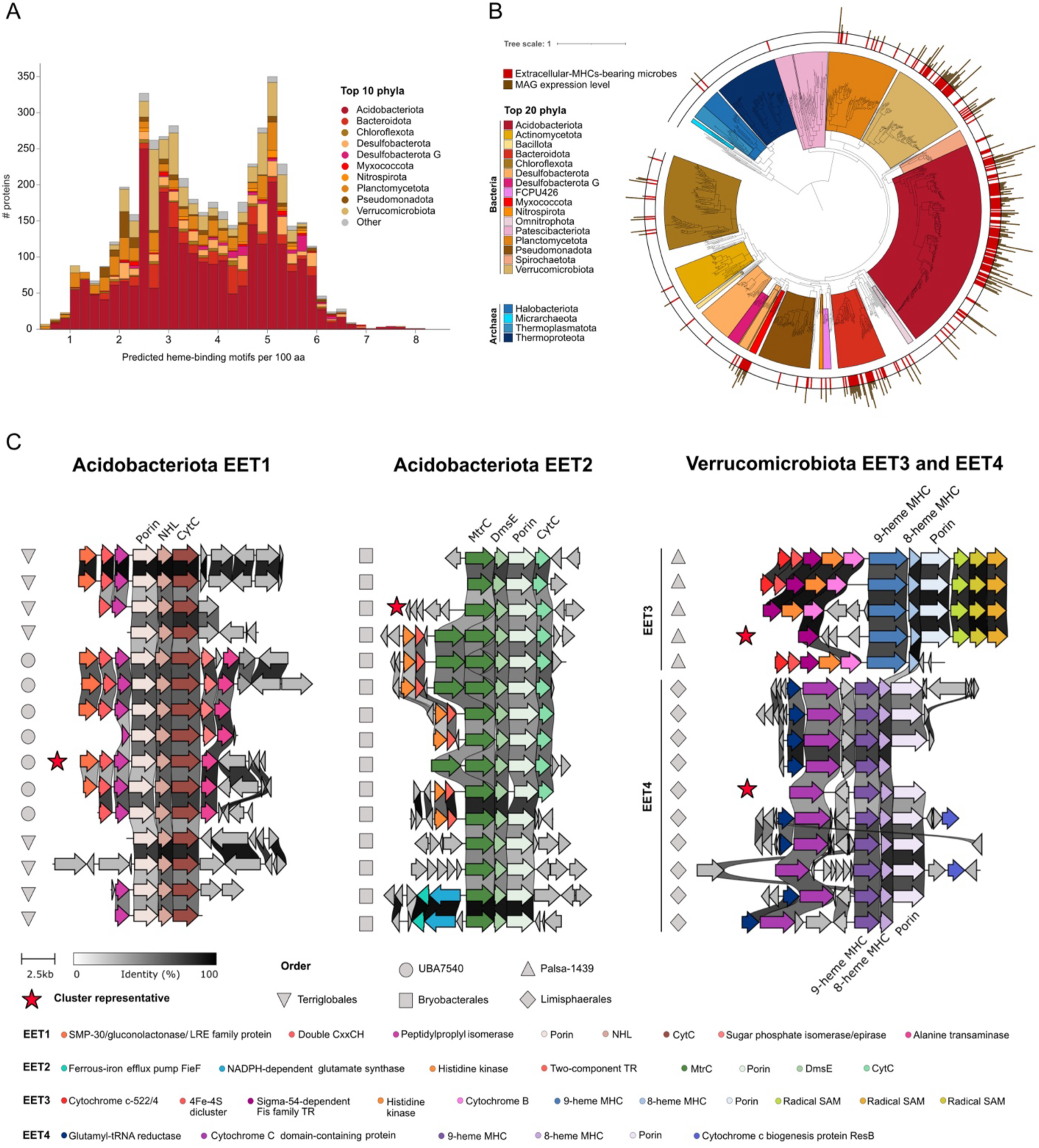
Multiheme cytochromes are widespread, diverse and transcriptionally active. **A**. Distribution of heme-binding motif density (motifs per 100 aa) across high-priority MHC proteins, stacked by the top 10 phyla. **B.** Species-level phylogeny of MAGs colored by the top 20 phyla. The outer red ring highlights MAGs encoding ≥5 extracellular MHCs and the external barplot reports the normalized expression level of MAGs carrying them across all samples. **C.** Conserved EET gene clusters identified in Acidobacteriota (EET1 and EET2) and Verrucomicrobiota (EET3 and EET4) across MAGs. Genes have different colors according to their functional annotation. Shapes indicate the taxonomic order of each cluster-bearing MAG and red stars mark the representative clusters. The bar below the alignments shows the amino acid identity (%) in each alignment.

High-priority MHC candidates were primarily associated with Bacteria. Despite Archaea representing 19% of the recovered MAGs, only ∼0.19% (9 proteins) of Tier 1 MHCs were associated with this domain. The vast majority of Tier 1 MHCs were encoded by Acidobacteriota (∼53%) and Verrucomicrobiota (∼13%). Strikingly, extracellular MHCs were concentrated in MAGs that were transcriptionally active across the sampled peat profiles (**Figure 3B**). Most of the 50 highest-expressed EET candidates across both shallow and deep layers were attributed to Acidobacteriota (32 MAGs, including 14 Terriglobia and 10 Bryobacterales) and Verrucomicrobiota (5 MAGs). This reveals that the diverse MHC collection is encoded by dominant and transcriptionally active peat bacteria.

### Acidobacteriota and Verrucomicrobiota have conserved and actively expressed clusters that could perform EET

Having identified putative EET-relevant MHCs in Acidobacteriota and Verrucomicrobiota, we next asked whether these proteins were embedded within broader genomic regions consistent with EET. For this, we extracted the genomic regions (±5 genes) containing extracellular MHC genes from all Acidobacteriota (156 contigs, 111 MAGs) and Verrucomicrobiota (54 contigs, 39 MAGs) MAGs and searched for co-localized outer membrane-spanning structural proteins (porins), outer-membrane (assembly) proteins and periplasmic electron-transfer components that are necessary to link intracellular redox metabolism to extracellular TEAs.

Aligning the genomic regions containing putative EET-facilitating MHCs revealed four dominant syntenic architectures, with two in each phylum (**Figure 3C**; **Supplementary discussion**). All four architectures encoded combinations of extracellular or outer-surface MHCs, periplasmic cytochromes and predicted outer-membrane porins, consistent with pathways capable of transferring electrons across the cell envelope. In Acidobacteriota, these included a porin-scaffold-cytochrome system (EET1) and an Mtr-like porin-cytochrome conduit in the Terriglobia order Bryobacterales (EET2). In Verrucomicrobiota, they comprised analogous porin-multiheme cytochrome assemblies associated primarily with Palsa-1439 (EET3) and Limisphaerales (EET4). Although these architectures resembled known EET systems in organization (**ED Figure 6**), representative proteins shared only low (∼8-31%) sequence identity with characterized Mtr/Omc components (**ED Figure 7**). This divergence was further supported by phylogenetic analysis, which revealed that porins and cytochromes from each architecture formed distinct clades that lacked close homologues at ≥50% sequence identity (**Figure 4A**). These findings support the identification of four conserved, sequence-divergent EET-like architectures in Acidobacteriota and Verrucomicrobiota that are organized similarly to canonical conduits but differ in amino acid composition and form distinct phylogenetic clades.

**Figure 4.**
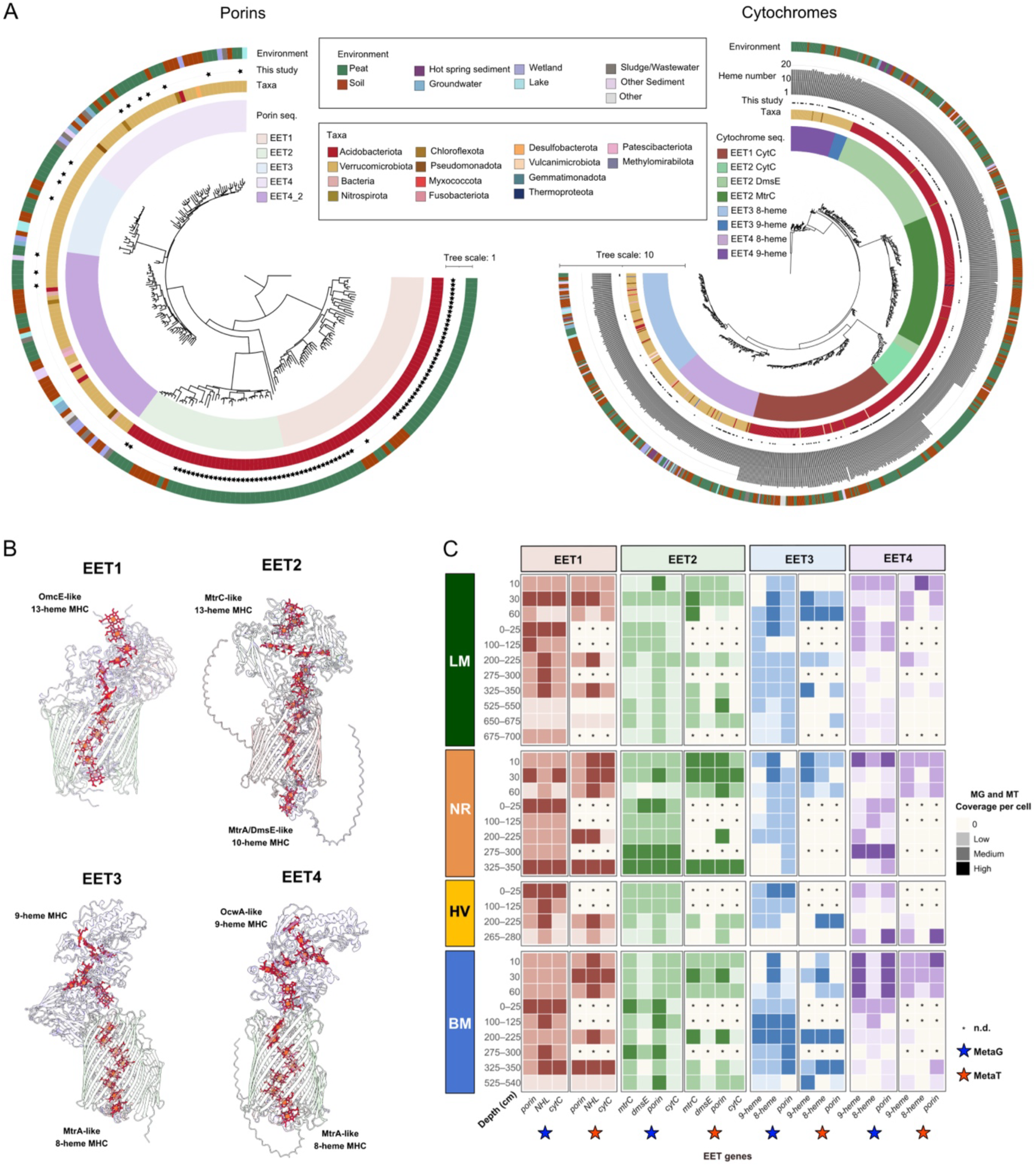
Members of the phyla Acidobacteriota and Verrucomicrobiota have conserved clusters likely involved in EET. **A.** Phylogenetic trees of conserved porins (left) and multiheme cytochromes (right) across EET systems and retrieved from publicly available databases. Sequences sharing ≥ 50% identity were retrieved prior to tree reconstruction. Colored outer rings indicate the environmental origin of each sequence, whereas black stars highlight sequences from this study. Inner rings indicate taxonomic affiliation (taxa = phyla), number of heme-binding motifs (CXXCH + CXXC_n_H + CX_n_CH, for cytochromes tree) and corresponding EET porins or cytochromes. **B.** Predicted structural models of representative proteins from each EET cluster. Heme iron atoms are highlighted in red. The EET clusters display distinct structures, including OmcE-like, MtrC-like, MtrA-like and OwcA-like MHCs with variable numbers of heme groups (8-13). **C.** Heatmap showing the mean normalized metagenomic and metatranscriptomic coverage (Coverage per cell: MG and MT coverage per cell) of the core EET clusters genes across each peatland and depth (cm). Expression levels were grouped into four categories (0, Low, Medium, High) using thresholds derived from the 25^th^ and 75^th^ percentiles of non-zero values across all clusters both for the metagenomic (blue star, metaG) and the metatranscriptomic (red star, metaT) datasets. n.d (no data) indicates absence of metatranscriptomic data for a specific sample depth. LM: Lungsmossen; NR: Norra Romyren; HV: Havsjömossen; BM: Björsmossen.

Further supporting their potential role in EET, structural modelling indicated that these sequence-divergent systems converge on architectures compatible with EET. Representative proteins showed structural similarity to Mtr/Omc-type components from *Shewanella*, *Geobacter* and *Thermincola*, and multimer modelling supported the assembly of membrane-spanning complexes resembling Mtr-like conduits (**Figure 4B**; **ED Figure 6**). In addition, the modelled architecture of all four EET systems contained heme-heme distances <14Å, which is consistent with efficient intra-protein electron hopping^21^ (**ED Figure 8**). Together, these structural features support the interpretation that the four divergent syntenic architectures encode functional multiheme electron-transfer conduits.

The core genes of all four architectures were actively expressed across peat profiles, including in deep layers (**Figure 4C**), indicating that these systems are active even where canonical inorganic electron acceptors are undetectable. Although the electron donors and acceptors for these microbes remain unresolved, our data suggest that deep peat may be redox-active, possibly sustaining EET *in situ*. Collectively, the syntenic conservation, structural convergence of Mtr-type architectures and expression across depth point toward a shared strategy employed by clades of Acidobacteriota and Verrucomicrobiota: EET via multiheme relays that could couple intracellular metabolism to extracellular electron acceptors, with peat POM representing one plausible candidate.

### Complex organic matter degradation and H_2_ cycling potentially fuel EET systems

Since conserved EET genes were expressed in both spatial and deep profiles (**Figure 4C**), we reconstructed the metabolism of EET-bearing Acidobacteriota (96 MAGs) and Verrucomicrobiota (24 MAGs) to identify the carbon and energy sources that could support these systems, especially under the more reducing conditions expected at depth (**Figure 5A,B; Supplementary discussion**). Metabolic reconstruction of these MAGs exhibited predominantly heterotrophic lifestyles with transcriptionally active CAZymes indicating the degradation of structurally complex, aromatic-rich organic matter into sugars that enter glycolysis and the Entner-Doudoroff pathway, generating pyruvate. This could then be oxidized to acetyl-CoA by the anaerobic pyruvate ferredoxin: oxidoreductase (PFOR), generating reduced ferredoxin (Fd_red_). Together with reducing equivalents produced by the TCA cycle, glycine cleavage system and peptide and protein degradation, this provides several routes by which organic carbon oxidation could generate electrons potentially feeding the respiratory chain and finally the EET systems.

**Figure 5.**
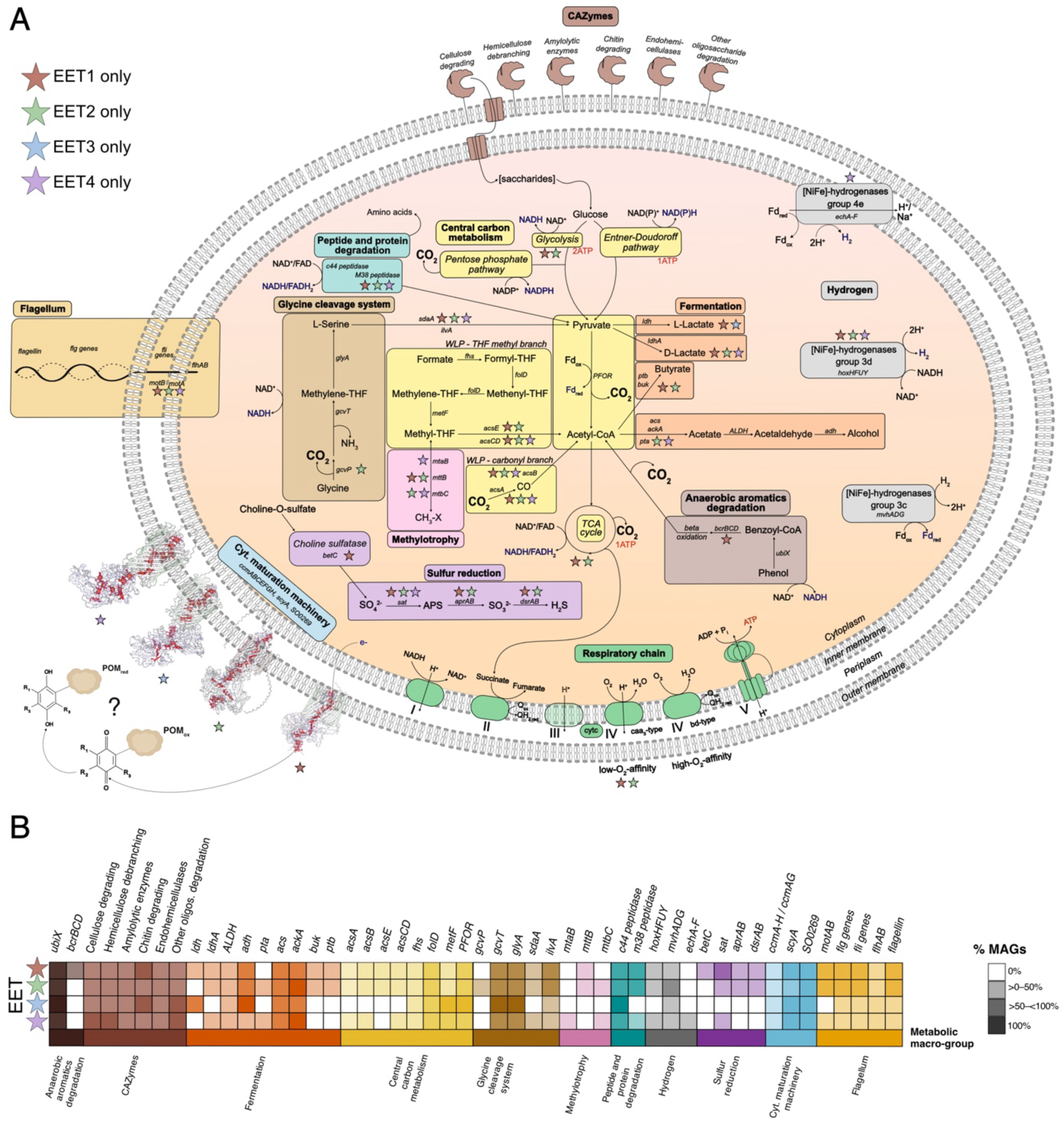
Metatrascriptome-inferred metabolic reconstruction of Acidobacteriota and Verrucomicrobiota encoding conserved EET machinery. **A.** Metabolic reconstruction of Acidobacteriota and Verrucomicrobiota MAGs carrying EET conserved machinery (EET1: Terriglobales, UBA7540; EET2: Bryobacterales; EET3: Palsa-1439; EET4: Limisphaerales). Metabolic potential is inferred from metagenomes and activity supported by gene expression in metatranscriptomes. For central carbon metabolism and respiratory-chain complexes, displayed pathways are both encoded in at least 50% of the MAGs and with ≥ 70% mean completeness and also with detected expression of more than 50% of pathway genes. All displayed functions are shared within Acidobacteriota (EET1 and EET2) and Verrucomicrobiota (EET3 and EET4) unless indicated by stars, which highlight pathways or genes fully encoded and transcribed in specific EET systems. Genes for each pathway or system are shown in italics. ATP-generating pathways are shown in red, while reduced electron carriers or electron donors potentially feeding EETs are shown in blue. **B.** The heatmap shows the percentage of MAGs encoding each EET machinery, in which each function is both encoded and transcribed. Functional categories are indicated by color and correspond to the colors used in the metabolic reconstruction. CAZymes: Carbohydrate-Active Enzymes; POM: particulate organic matter; WLP: Wood-Ljungdahl pathway; Cyt. maturation machinery: Cytochrome *c* maturation machinery.

Hydrogen may provide additional reducing power, especially in deep peat. EET-bearing MAGs expressed various [NiFe]-hydrogenases genes. These included group 3c hydrogenases (*mvhADG*), which oxidize H_2_ and generate Fd_red_. In turn, group 3d (*hoxHFUY*) hydrogenases may contribute to H_2_ production or consumption, depending on local redox conditions. Further, group 4e (*echA-F*, EET4 only) hydrogenases, which couple reduced ferredoxin oxidation to H_2_ production, may function as energy-conserving hydrogenases^22^. This pattern suggests active H_2_ cycling and points to H_2_ as a potential electron donor supporting microbial persistence and survival in low-energy deep layers^23^. All EET-bearing lineages also expressed high-O_2_-affinity *bd*-type terminal oxidases. This indicates a capacity to exploit locally available O_2_ as TEA, including in microoxic microniches where O_2_ is transiently supplied by water-table fluctuations, plant aerenchyma^1^ or co-occurring aerobic metabolisms. Acidobacteriota also expressed low-O_2_-affinity *caa*3-type oxidases, suggesting greater flexibility to adjust respiration across a broader range of O_2_ availabilities. Expression of cytochrome *c* maturation (*ccm*) machinery genes^24^, necessary for MHC assembly, further supports the functional use of EET systems *in situ*. Together, these results suggest that EET-bearing Acidobacteriota and Verrucomicrobiota primarily use structurally complex organic matter as carbon and electron source, with H_2_ as an additional electron donor, respire O_2_ when available, and employ their EET complexes to redirect electrons to an extracellular TEA when soluble TEAs become limiting.

### EET-bearing MAGs and gene clusters are distributed across other ecosystems

To gain insights into the ecological relevance of these EET systems, we searched for the conserved core genes in metagenomes and MAGs beyond the Värmland bogs. Comparisons to SPRUCE and Stordalen Mire revealed near-identical matches in 29 MAGs, spanning Terriglobales, UBA7540, Bryobacterales, Palsa-1439 and Limisphaerales. Broader searches using Sandpiper^25^ and the mOTUs database^26^ showed that EET-bearing taxa were not confined to peatlands but are distributed across diverse ecosystems, including seawater, freshwater lakes, glacier-fed stream sediment, forest soils and hot springs. Across these environments, we identified 184 MAGs from 103 species encoding at least one EET-associated gene, predominantly affiliated with Acidobacteriota and Verrucomicrobiota. Applying a stricter threshold requiring at least three co-occurring genes from the same EET cluster architecture, indicative of a more complete transfer complex, yielded 119 MAGs from 66 species, demonstrating that EET machinery is widespread well beyond the ecosystems in which it was first discovered (**Supplementary Table 1**).

Together, these analyses indicate that Acidobacteriota and Verrucomicrobiota EET systems are conserved across geographically distant peatlands and occur more broadly across anoxic or redox-stratified environments, supporting MHC-mediated EET as a potentially widespread but overlooked metabolic strategy among members of these lineages.

## Discussion

Through depth-resolved metagenomics and metatranscriptomics across deep peat profiles in four Swedish ombrotrophic bogs, we provide ecological, genomic and transcriptional evidence for previously unrecognized EET-related respiratory capacity in these ecosystems. First, we show that microbial communities spanning Acidobacteriota, Verrucomicrobiota and diverse anaerobic lineages remain structured and transcriptionally active from the near-surface acrotelm through the deep catotelm, with depth as the primary axis of compositional change. Second, Acidobacteriota and Verrucomicrobiota encode exceptionally large, sequence-divergent MHC repertoires organized into conserved porin-MHC gene clusters with structural features compatible with EET. Third, these systems are transcriptionally active across all sampled depths, including the deep catotelm where canonical inorganic TEAs are scarce. Together, these findings reveal that EET-capable microbial machinery is a recurrent and actively expressed feature of dominant uncultured peat bacteria, with implications for how anaerobic carbon cycling and methane emissions from northern peatlands are understood and modelled.

A central finding of this study is the identification of conserved, transcriptionally active and structurally supported multiheme cytochrome systems in Acidobacteriota and Verrucomicrobiota, two dominant but comparatively understudied peatland lineages. EET has been most extensively characterized in metal-reducing bacteria such as *Geobacter sulfurreducens* and *Shewanella oneidensis*, where Mtr/Omc-like systems connect intracellular metabolism to extracellular minerals or soluble electron shuttles^11^. By contrast, the EET-like architectures identified here are deeply sequence-divergent from canonical systems yet retain key organizational and structural features compatible with electron transfer across the cell envelope. Their expression across shallow and deep peat profiles, together with the near-absence of dissolved Fe(III) and Mn(IV) as potential TEAs, suggests that these systems may not primarily target inorganic metal oxides, pointing instead towards peat organic or as-yet-unidentified extracellular electron acceptors. Building on earlier genomic indications of porin-MHC gene clusters in Acidobacteriota and freshwater systems^13,14^, our study provides the first characterization of these systems at the level of genomic architecture, structural modelling, phylogenetic placement and transcriptional activity across depth. Thus, our study identifies the microbial lineages and candidate EET complexes most likely to mediate EET in ombrotrophic peat, providing a genomic and transcriptomic framework for mechanistic follow-up.

These findings have implications for how carbon flow is conceptualized in peatlands. Rather than a simple progression from aerobic respiration near the surface to fermentation and methanogenesis at depth, our data suggest a more distributed redox network in which complex organic matter degradation, hydrogen cycling, transient oxygen use and EET jointly sustain microbial metabolism. In this framework, redox-active organic matter may act not only as a carbon reservoir but also as an electron-accepting matrix. Specifically, POM quinone moieties can accept electrons from MHC conduits, acting as the TEA for EET. Upon reoxidation, POM donates those electrons to a downstream oxidant. As long as reoxidation is maintained, POM functions not as a consumed terminal sink but as a regenerable redox shuttle cycling between reduced and oxidized states. In the upper peat layers (0-60 cm), transient oxygen inputs through water table fluctuations, root aerenchyma or co-occurring aerobic metabolisms could provide localized microoxic conditions for POM reoxidation. In the deep catotelm (>200 cm), where conditions are strictly anoxic, however, POM reoxidation, potentially through hypothetical as-yet uncharacterized anaerobic oxidative processes, remains a critical unresolved question. While cable bacteria (*Electrothrix, Electronema*) perform centimeter-scale long-range electron transfer and provide a useful mechanistic parallel for how electrons can be transported over distance, they have so far primarily been described in aquatic sediments^12^ rather than peatlands. Notably, recent work has demonstrated that the freshwater cable bacterium *Electronema aureum* GS can sustain anoxic growth via EET to insoluble electron acceptors at rates comparable to aerobic respiration, mediated by outer-membrane cytochromes and riboflavin electron shuttles^27^. This underscores the energetic competitiveness of EET-based respiration via MHCs and broadens the ecological contexts in which it may operate. In ombrotrophic bogs, a comparable mechanism could provide a route for reoxidizing reduced POM under otherwise anoxic conditions. Whether EET machinery across depth gradients in Acidobacteriota and Verrucomicrobiota could contribute to such longer-range electron transfer through syntrophic cell networks, soluble electron shuttles or a conductive peat matrix remains unresolved, and analogous processes may yet be found in other uncharacterized peatland lineages.

Looking forward, the identification of widespread, actively expressed EET machinery opens a new phase of investigation into anaerobic carbon cycling in peatlands. It should first be noted that dissolved CO_2_:CH_4_ concentration ratios may partly reflect differential gas bubble partitioning at depth, since CH_4_ is less soluble than CO_2_ (Henry’s law), potentially causing dissolved ratios to overestimate the true respiratory stoichiometry. Quantifying gas bubble volumes alongside porewater profiles would allow this effect to be resolved. Beyond this, critical open questions remain: what is the identity of the electron acceptor(s) in the deep catotelm, where inorganic TEAs are undetectable and POM reoxidation is uncertain? Could dissolved organic matter (DOM) act as a mobile electron shuttle, reduced by MHC conduits and transferring electrons to a secondary, as-yet-unidentified extracellular acceptor? Are there depth-dependent shifts in the identity of the terminal acceptor as peat chemistry changes with age and recalcitrance? Addressing these questions will require combining the genomic and transcriptomic framework established here with direct electrochemical measurements, isotope-labelling approaches, and enrichment or cultivation of the uncultured lineages encoding these systems.

Our findings reveal previously unrecognized EET machinery in dominant, uncultured peat bacteria, providing the first candidate molecular framework for the long-observed discrepancy between CO_2_ and CH_4_ production in ombrotrophic northern peatlands and identify Acidobacteriota and Verrucomicrobiota as the lineages most likely to mediate it. Beyond peatlands, the convergent evolution of sequence-divergent EET architectures in these abundant environmental phyla suggests that EET may represent a widespread respiratory strategy across anoxic terrestrial and aquatic environments. Incorporating this process into peatland biogeochemical models may therefore be necessary to accurately represent anaerobic carbon cycling, and ultimately the methane emissions, from one of Earth’s largest terrestrial carbon stores.

## Methods

### Field sites and sampling

Four pristine ombrotrophic bogs in the Värmland County, central Sweden, were sampled in June 2024: Björsmossen (BM; 59.69276°N, 014.27728°E), Norra Romyren (NR; 59.86159°N,014.83193°E), Havsjömossen (HV; 59.87202°N, 014.85343°E) and Lungsmossen (LM; 59.54800°N,014.23829°E) (**Supplementary Methods**).

**Spatial profiling** was carried out in BM, NR, and LM across three sites (site 1-3) located a few meters apart. Peat samples were taken in 10 cm intervals up to 60 cm depth and distributed in sterile Whirl-Pak bags and frozen immediately on dry ice or directly put into sealed glass jars.

**Deep profiling** was performed using a Russian peat sampler (Royal Eijkelkamp, Giesbeek, The Netherlands), collecting 50 cm soil sections and reaching maximum depths of 540 cm (BM), 350 cm (NR), 700 cm (LM) and 280 cm (HV). Each 50 cm segment was divided into two equal halves (e.g., 0-25 cm, 25-50 cm), stored in Whirl-Pak bags, and frozen on dry ice. Samples were kept frozen during transport and stored at 4°C (jars) and −80°C (Whirl-Paks) until further processing.

Samples used for 16S rRNA gene amplicon sequencing are listed in **Supplementary Table 2**, while those used for metagenomic and metatranscriptomic sequencing are listed in **Supplementary Table 3**.

### Biogeochemical analyses

*In situ* pH, conductivity, and temperature were measured using a calibrated WTW Multi 430i instrument (Xylem Analytics, Germany) at BM, LM and NR during spatial profiling. For dissolved gas measurements, 8 ml porewater was injected into N_2_-flushed 20 ml glass vials containing 3 g NaCl to promote gas transfer to the headspace. CO_2_ and CH_4_ were quantified suing a GC-Methanizer system (SRI 8610C GC, SRI Instrument Europe, Bad Honnef, Germany) operating at 40° C and equipped with Porapak Q and HayeSep D columns connected in series. Nitrogen (N_2,_ grade 5.0) was used as carrier gas and Hydrogen (H_2_, grade 5.0, all gases from Linde Gas) as the fuel for the methanizer detector (a modified flame-ionization detector). CH₄ was quantified directly, whereas CO₂ was first reduced to CH₄ by H₂ on a nickel catalyst operated at 380 °C to enable its detection and quantification.

Peat collected in jars from BM and NR was processed under strictly anoxic conditions inside an N_2_-filled glove box (<2.3 ppm O_2;_ Mbraun, Germany). Porewater was extracted by squeezing, filtered (0.45 µm, Sarstedt, Germany), and analyzed for dissolved species. Major anions (SO_4_^2−^, NO_3_^−^) were quantified by ion chromatography (IC) (Metrohm 940 Professional IC Vario, A Supp 5, 250/4.0 column), and cations (total Fe and total Mn) were quantified by inductively coupled plasma-mass spectrometry (ICP-MS) using an Agilent 7900 instrument (Basel, Switzerland). Organic acids were detected using High-Performance Liquid Chromatography (HPLC) (Agilent 1260 Infinity II) with a Hamilton PRP X-300 column (7µm, 4.1 x 250 mm).

### Nucleic acid extractions and Illumina sequencing

For 16S rRNA gene amplicon sequencing, DNA was extracted using the DNeasy PowerSoil Pro Kit (Qiagen, Germany) following the manufacturer’s instructions. For metagenomics and metatranscriptomics, total DNA and RNA were co-extracted using the RNeasy PowerSoil Total RNA Kit, followed by DNA recovery from the same lysate with the RNeasy PowerSoil DNA Elution kit (Qiagen, Germany). Nucleic acid purity and concentration were assessed using the NanoDrop2000 and Qubit 4 fluorometer (ThermoFisher Scientific, USA).

Sequencing was performed by Novogene (Novogene, Ltd, Cambridge, UK) on an Illumina NovaSeq 6000 platform (paired-end 250bp) for 16S rRNA amplicons, and on an Illumina NovaSeq X Plus platform (paired-end 150bp) for metagenomic and metatranscriptomic libraries.

### 16S rRNA gene amplicon sequencing and statistical analyses

The V4 region of the 16S rRNA gene was amplified using primers 515F (5’-GTGCCAGCMGCCGCGGTAA-3’) and 806R (5’- GGACTACHVGGGTWTCTAAT-3’), targeting both archaeal and bacterial diversity^28^. Raw paired-end 16S rRNA gene reads were quality-checked with FastQC (v0.12.01)^29^ and PhiX contamination was removed using Bowtie2 (v2.5.4)^30^. Trimming of adapters and primers was performed via Cutadapt (v4.9)^31^. Paired-end read merging, Amplicon Sequence Variant (ASV) inference (UNOISE3) and *de novo* chimera removal (UCHIME3) were conducted with vsearch (v2.22.1)^32^, after which ASVs were clustered into 97% OTUs.

Taxonomic classification of ASVs was performed using the Silva v138 database^33^. Alpha and beta diversity analyses were conducted in R (v4.5.2) using the *vegan*^34^ and *ggplot2*^35^ packages. The OTU count tables were rarified to the minimum sequencing depth over 100 iterations. Observed richness and Shannon diversity were calculated for each rarefied table and averaged across iterations, while Pielou’s evenness was derived from the averaged Shannon diversity and richness estimates. Alpha diversity metrics were compared across samples using two-way ANOVA on best Normalized-transformed values. Beta diversity was assessed using Bray-Curtis and Jaccard dissimilarities and Bray-Curtis dissimilarities were also visualized by distance-based redundancy analysis (dbRDA). Relationships between microbial community composition and environmental variables were evaluated with PERMANOVA (adonis2, 999 permutations) and envfit (999 permutations).

### Metagenomic and metatranscriptomic analyses

For metagenomics, analysis involved quality assessment of the raw reads with fastQC (v0.12.1)^29^ and trimming with BBDuk (BBMap v39.01)^36^ (ktrim=r, k=23, 106 mink=11, hdist=1, trimpolyg=20, trimq=20, qtrim=rl, minlen=105, tpe, tpo). Trimmed reads underwent *de novo* assembly using metaSPAdes (v4.0.0)^37^ (k-mers: 21,33,55,77,99 and 127) and resulting assemblies were quality-checked using SeqKit (v0.10.1)^38^ and BBMap (v.39.01)^36^, discarding scaffolds shorter than 1000 bp (**Supplementary Table 4**).

Open reading frames (ORFs) were predicted from assembled scaffolds using Prodigal (--mode anon) (v2.11.0)^39^ Binning was performed via MetaBAT2 (v2.17)^40^, CONCOCT (v1.1.0)^41^, and Maxbin2 (v2.2.7)^42^. The best bins were selected using DAS Tool (v1.1.7)^43^ and metagenome-assembled genomes (MAGs) at species level (average nucleotide identity threshold (ANI) ≥95%) were obtained via dereplication with dRep (v3.4.5)^44^. Completeness and contamination of MAGs were estimated via CheckM (v1.2.2)^45^ retaining only medium- to-high-quality MAGs according to the MIMAG standards^46^. For their taxonomic classification, GTDB-Tk (v2.4.0)^16^ was employed, using the GTDB r220 database. To identify candidate novel species, GTDB-Tk ani_rep was run on the 1081 species level dereplicated MAGs, extracting ANI and alignment fraction (AF) to the closest GTDB representative genome (ani_closest). High-quality MAGs (>90% completeness, <5% contamination) with ANI below the species ANI circumscription radius of the closest GTDB reference genome (typically 95%) and/or AF below the GTDB assignment threshold (65%)^47^ were considered candidate novel species.

The relative abundance of each MAG across samples was determined with CoverM (v0.6.1)^48^, by mapping the trimmed reads back to the dereplicated genomes (**Supplementary Table 5**). Functional annotations and genome-level metabolic reconstruction were performed with DRAM (v1.5.0)^49^ and METABOLIC-G (v4.0)^50^. InterProScan (v101.0)^51^ was used as an additional functional annotation tool.

Microbial community composition was also inferred using raw metagenomic reads using SingleM^25^.

To compare our Värmland metagenomes with Stordalen Mire and SPRUCE sites, MASH distances (v2.1)^52^ were calculated at the metagenomic read level. 32 metagenomes per site were selected and identified via the SRA Run Selector (https://www.ncbi.nlm.nih.gov/sra, see **Data Availability**). MASH sketches were generated using mash sketch -s 10000 -r -m 2 and outcome visualized by Non-metric MultiDimensional Scaling (NMDS) plot on R using the *vegan*^34^ and *ggplot*^35^ packages.

Metatranscriptomic raw reads were processed using TranscriptM (v0.5)^53^ (including read trimming via Trimmomatic (v0.32)^54^, quality assessment via FastQC (v0.10.1)^29^, PhiX removal using fxtract (v1.2) (https://github.com/ctSkennerton/fxtract) and rRNA, tRNA, tmRNA depletion with SortMeRNA (v2.0)^55^. Cleaned reads were aligned to the MAGs using Bamm (v.1.5.0), (https://ecogenomics.github.io/BamM/) and gene-level coverage and read counts were calculated using GFF annotations provided for each genome.

### Metatranscriptome normalization

#### Construction of a non-redundant, functionally annotated gene catalogue

To facilitate comparisons on the genetic content of microbial communities across samples, we constructed a non-redundant gene catalogue (see **Data availability**). Genes were predicted from all metagenomic contigs using pyrodigal-GV (v0.3.2)^56^ and subsequently clustered at 95% nucleotide identity with a minimum of 90% horizontal coverage using the easy-linclust algorithm of MMSeqs2 (v18.8cc5c; parameters: --min-seq-id 0.95 -c 0.9)^57^. The gene cluster representatives were retained to form a non-redundant gene catalogue. The gene catalogue was functionally annotated using HMMER (hmmsearch) (v3.4)^58^, with the gathering cut-off threshold (parameter: --cut_ga), against the KEGG orthology (obtained 03.2025)^59^, Pfam (v37.0)^60^ and METABOLIC^50^ HMM databases. The best-scoring hit for each gene cluster, determined based on the bitscore and e-value, was retained.

#### Estimating the abundance, transcription and expression of genes, protein families and MAGs

To estimate the abundance and transcription of gene clusters, we profiled the gene catalogue across all metagenomes and metatranscriptomes through read alignments with the bwa mem tool (v0.7.19; parameters: -a)^61^. The aligned reads were filtered to retain those with an alignment nucleotide identity of ≥95%. From the retained read alignments, we calculated the horizontal and vertical coverage. Vertical coverage (i.e., mean depth per base) was calculated as the total number of aligned bases divided by the gene length, scaled to 1 kilobase. Genes with a horizontal coverage <95% were removed. To account for differences in sequencing depth across samples, we estimated the number of genomes sequenced by taking the median of the summed vertical coverage of 40 single-copy marker genes per sample^62^. The vertical coverage of each gene cluster was subsequently normalized by the number of genomes sequenced to generate copies per genome.

To infer the expression of each gene in samples where both metagenomes and metatranscriptomes were generated, we normalized the copies per genome from metatranscriptomes by the copies per genome from the metagenome before log2 scaling. The computed expression thus represents the estimated expression per gene in relation to the number of copies of that gene. That is, an expression value of 0 would indicate that the coverage in the metatranscriptome is equal to the coverage in the metagenome, and thus, on average across the population, there is no up or down regulation. In contrast, a value greater than 1, indicates a transcription value that is higher than expected based on the number of copies of the gene in the population, suggesting an up-regulation.

To estimate the coverage and expression of protein families, we leveraged coverage information of each gene cluster and their annotations against the KEGG^59^ and Pfam databases^60^. The copies per genome of each protein family was determined as the sum of the respective gene clusters. The expression of each protein family was estimated following the same processes as outlined above.

To estimate the coverage and expression of MAGs, we used information from the 40 single-copy marker genes. For each MAG, we calculated the median copies per genome across its encoded 40 single-copy marker genes and normalized this value by the number of genomes sequenced, resulting in an estimate of the proportion of genomes sequenced that are attributed to that MAG. The expression of each MAG was then estimated following the same process as outlined above.

### Detection of multiheme cytochromes and protein family clustering

Putative multiheme cytochromes (MHCs) were identified from all predicted protein sequences using custom Python (v3.11.6) scripts. Proteins containing ≥ 3 heme-binding motifs (CXXCH, CXXC_n_H, or CX_n_CH, with up to 70 residues between C and H) of the same type were retained as candidate MHCs. Each MHC was linked to its corresponding MAG or contig, distinguishing binned and unbinned fractions. Functional annotations were integrated from DRAM^49^ and InterProScan^51^ and expression levels of the respective genes were retrieved from TranscriptM^53^ analysis (metatranscriptomic analysis). Subcellular localization of proteins was predicted using PSORTb (v3.0) (--negative, since the majority of the MHC-bearing microbes identified in the study are Gram-negative bacteria)^63^.

To investigate potential patterns of sequence similarity and functional conservation among the putative MHCs, sequence-level protein clustering was performed using MMseqs2 (v14-7e284; parameters: --min-seq-id 0.50 -c 0.9)^57^. This approach allowed to group homologous proteins and decrease redundancy across the entire dataset, while preserving representative diversity. As a reference, 73 and 23 MHCs protein sequences from *Geobacter sulfurreducens* PCA Δ*omcS* strain^19^ and *Shewanella oneidensis* MR-1 strain^20^, respectively, were downloaded from UniProt (https://www.uniprot.org/) and included in the clustering workflow.

To assess whether microorganisms carrying MHCs are phylogenetically related, we constructed a phylogenetic tree from the 1081 dereplicated MAGs (species level) using GToTree (v1.8.6)^64^ (marker set: Bacteria_and_Archaea.hmm), visualized in iTol (v7.3)^65^ and rooted at the Bacteria-Archaea split. MAGs carrying ≥5 extracellular MHCs have been selected for decorating the tree and their expression level across all samples comes from normalized metatranscriptomic data.

### Multiheme cytochrome credibility scoring

Candidate MHC sequences were first clustered by SeqC_ID and the credibility of each cluster was assessed from the combined properties of its member proteins using a custom Python script (see **Data availability**). The score was based on the proportion of members with ≥6 or ≥10 heme-binding motifs, the median number of motifs per cluster, PSORT^63^ localization, PFAM^60^ annotation, heme density and metatranscriptomic expression support. Localization supported clusters enriched in periplasmic, outer-membrane or extracellular predictions, with cytoplasmic-membrane predictions considered weaker support and cytoplasmic predictions penalized. PFAM annotations were summarized as the fraction of cluster members carrying curated cytochrome c/multiheme PFAMs, outer-membrane EET-related PFAMs or PFAMs considered negative evidence. Core cytochrome *c*/multiheme support included PF00034, PF02335, PF13435, PF14537, PF11783, PF02085, PF14522, PF07635, PF22678, PF09698, PF09699, PF22112 and PF22113, while outer-membrane EET support was assigned from PF22112, PF22113 and PF03264. Positive scores were assigned when ≥50% of members carried core cytochrome c/multiheme PFAMs or ≥20% carried outer-membrane EET PFAMs. Clusters were penalized when ≥50% of members carried PFAMs associated with non-multiheme proteins, including Fe–S proteins, molybdopterin oxidoreductases, radical SAM enzymes, ABC/UvrA-like proteins or tRNA-synthetase-like proteins. Heme density was calculated as the number of motifs per 100 amino acids and expression was included only as a minor support criterion using the maximum member-level expression within each cluster. Final scores classified clusters as Tier 1 credible candidates, Tier 2 putative candidates or tier 3 uncertain candidates, corresponding to scores of ≥6, 3-5 and ≤2, respectively.

### Gene cluster alignments

Genomic regions (±5 genes) surrounding extracellular MHC genes were extracted from all Acidobacteriota and Verrucomicrobiota MAGs (see **Data Availability**), annotated with Bakta^66^ (https://bakta.computational.bio/) and aligned using clinker (v0.0.32)^67^. Loci were grouped into conserved cluster types based on shared gene content and syntenic arrangement, and representative loci were selected for structural modeling. To compare these conserved cluster types with characterized EET systems, equivalent genomic regions were extracted around genes involved in EET from representative model organisms, including *Shewanella baltica* OS678^68^, *Shewanella oneidensis* MR-1^20^, *Geobacter sulfurreducens* PCA^19^, *Thermincola potens* JR^69^.

### Phylogenetic tree construction of MHCs and porins

Protein sequences were queried against the clustered non-redundant database (nr_clust) using BLAST^70^ to identify homologs. Retrieved sequences were filtered at ≥50% sequence identity and ≥90% alignment coverage prior to phylogenetic reconstruction. The selected sequences were aligned using MAFFT (v7)^71^ with the L-INS-i algorithm (--localpair --maxiterate 1000), and poorly aligned regions were removed with trimAl^72^ (-automated1). Maximum-likelihood phylogenetic trees were inferred using IQ-TREE3 (v3.0.1)^73^ with the best-fit substitution model determined by ModelFinder (-m MFP). Branch support was assessed with 1,000 ultrafast bootstrap replicates (-bb 1000).

### Protein modeling and structural analyses

For each conserved cluster representative, signal peptides were predicted using SignalP^74^ and removed prior to modeling. Monomeric structures of each representative conserved cluster were predicted using the AlphaFold3 server (https://alphafoldserver.com/). For MHCs predicted to span the outer membrane pore, the number of heme-binding motifs (canonical and non-canonical) determined the number of heme *c* ligands included in the prediction. Core EET cassette proteins were additionally modeled as heteromultimers and model confidence was evaluated by ipTM and pTM scores. Structural homologs were identified by searching against the PDB100 database using foldseek^75^. Predicted multimer models were superimposed onto experimentally determined structures from PDB using the Mathmaker tool in UCSF ChimeraX (v1.11)^76^. Hemes were labeled consecutively in order of their heme-binding motif in the polypeptide chain (N to C).

### Prevalence of conserved gene clusters across other environments

To test whether the genes coming from the conserved clusters identified in our sites are also present in other well-studied peatland systems, we searched for them in MAGs from NCBI BioProjects PRJNA1084886 (SPRUCE) and PRJNA888099 (Stordalen Mire). MAGs were screened using BLASTn-plus (v2.14.1)^70^ using nucleotide query sequences extracted from our reference genes. Hits were filtered (e-value ≤ 1×10⁻¹⁰, identity ≥ 95%, query coverage ≥ 95%, gapopen = 0), and a “credible” match required ≥1 essential EET gene plus ≥1 additional target gene on the same contig. NCBI MAGs were taxonomically classified with GTDB-Tk (v.2.4.0)^16^.

The dominant taxonomic classes identified in our dataset were further searched across other environments using Sandpiper^25^ (https://sandpiper.qut.edu.au/, accessed on 03.02.2026). We additionally screened all genomes in the mOTUs (v4.0.4)^26,77^ database for homologues of the identified EET gene sets. Matches were considered significant when they showed ≥50% sequence identity and ≥80% horizontal coverage.

### Acidobacteriota and Verrucomicrobiota metabolic reconstructions

Metabolic reconstructions were generated for Acidobacteriota and Verrucomicrobiota MAGs encoding conserved EET systems by integrating annotations from DRAM^49^ and METABOLIC^50^ outputs. Central carbon metabolism pathways were drawn in the reconstruction only if encoded in ≥ 50% of MAGs and with mean completeness of genes ≥ 70% (DRAM), and if more than 50% of the genes belonging to a pathway/module were found to be expressed according to the normalized metaT data (glycolysis, pentose phosphate pathway, Entner-Doudoroff pathway, TCA cycle, respiratory-chain complexes). WLP genes were retrieved from MAG annotations, while CAZyme annotations were retrieved from METABOLIC and their expression was also assessed according to the normalized metaT analysis. Other genes or pathways belonging to other metabolic functions were considered only when both encoded and transcribed in each EET system-bearing MAGs. For multi-subunit gene complexes, presence and transcription were shown only when the required subunits were both encoded and transcribed. For cytochrome *c* maturation machinery, only *ccmABCEFGH* (*ccmA-H*) genes were considered for EET1- and EET2-bearing Acidobacteriota, since *ccmD* was always not encoded by MAGs, whereas only *ccmA* and *ccmG* were considered from EET3- and EET4-bearing Verrucomicrobiota, as all remaining *ccm* genes were absent from MAGs in these systems.

## Supporting information

Fiorito_et_al_2026_Extended_Data_Figures

Fiorito_et_al_2026_Supplementary_information

## Data availability

16S rRNA gene amplicon, metagenomic and metatranscriptomic sequencing reads and MAGs reported in this paper are available under NCBI Bio-Project: PRJNA1468976. Reference datasets comprising Stordalen Mire metagenome datasets were selected from BioProjects PRJNA888099 PRJNA618049-PRJNA618067, PRJNA617210-PRJNA617211, PRJNA539550, PRJNA539561, PRJNA539566, PRJNA539572, PRJNA539574-PRJNA539576, PRJNA539579-PRJNA539580 and PRJNA518543-PRJNA518544. Reference SPRUCE metagenomes were selected from BioProjects PRJNA1084886, PRJNA444884-PRJNA444889 and PRJNA443591-PRJNA443617. Structural models, Genbank files of Acidobacteriota and Verrucomicrobiota contigs carrying extracellular MHCs, custom python script for MHC credibility scoring, all phylogenetic trees and fasta files used for their creation are available through Zenodo (https://doi.org/10.5281/zenodo.20393101). Scripts for generating the gene catalog from metagenomic datasets and performing gene expression analysis are available in Github (https://github.com/tpriest0/Profiling_metagenomes_and_mags). Additional Information is provided in Extended Data Figures 1-8 and Supplementary Information PDF and Supplementary Tables 1-5.

## Acknowledgements

This research was funded by the Swiss National Science Foundation (SNF 100001203), and a NOMIS Fellowship (awarded to TP). We thank Donat Crippa and Astrid von Mentzer for invaluable assistance with sampling and transport. We thank the Genetic Diversity Centre (GDC), ETH Zurich, for providing access to their facility and supporting some of the molecular work performed as part of the project. We thank Madison Barney and Björn Studer for help with the HPLC, IC and ICP-MS measurements.

## Author Contributions

GF, MS, MHS and MCS designed the study and collected the samples. DNA and RNA extractions were performed by GF. MHS performed porewater analyses to quantify dissolved gases. Amplicon and statistical analyses were performed by GF. Metagenomic datasets were constructed by GF adapting a pipeline developed by HZ. Metatranscriptomic datasets were constructed by GF adapting a pipeline developed by BD. Metatranscriptome normalizations were performed by TP. Multiheme cytochrome detection and protein family clustering were performed by GF. Genome alignments were performed by GF and MCS. MHC and porin phylogenetic trees were performed by BD. Structural predictions and searches were performed by MCS. Searching for MAG and EET module occurrence across datasets was performed by GF and TP. Metabolic reconstructions were performed by GF. The project was supervised by MCS with support from MS and MHS. Funding was acquired by MCS. GF, TP and MCS wrote the manuscript, with input from all coauthors.

