## Supplementary material for "Conserved multiheme cytochrome machinery for extracellular electron transfer is widespread and transcriptionally active across deep peat profiles": Fiorito_et_al_2026_Extended_Data_Figures

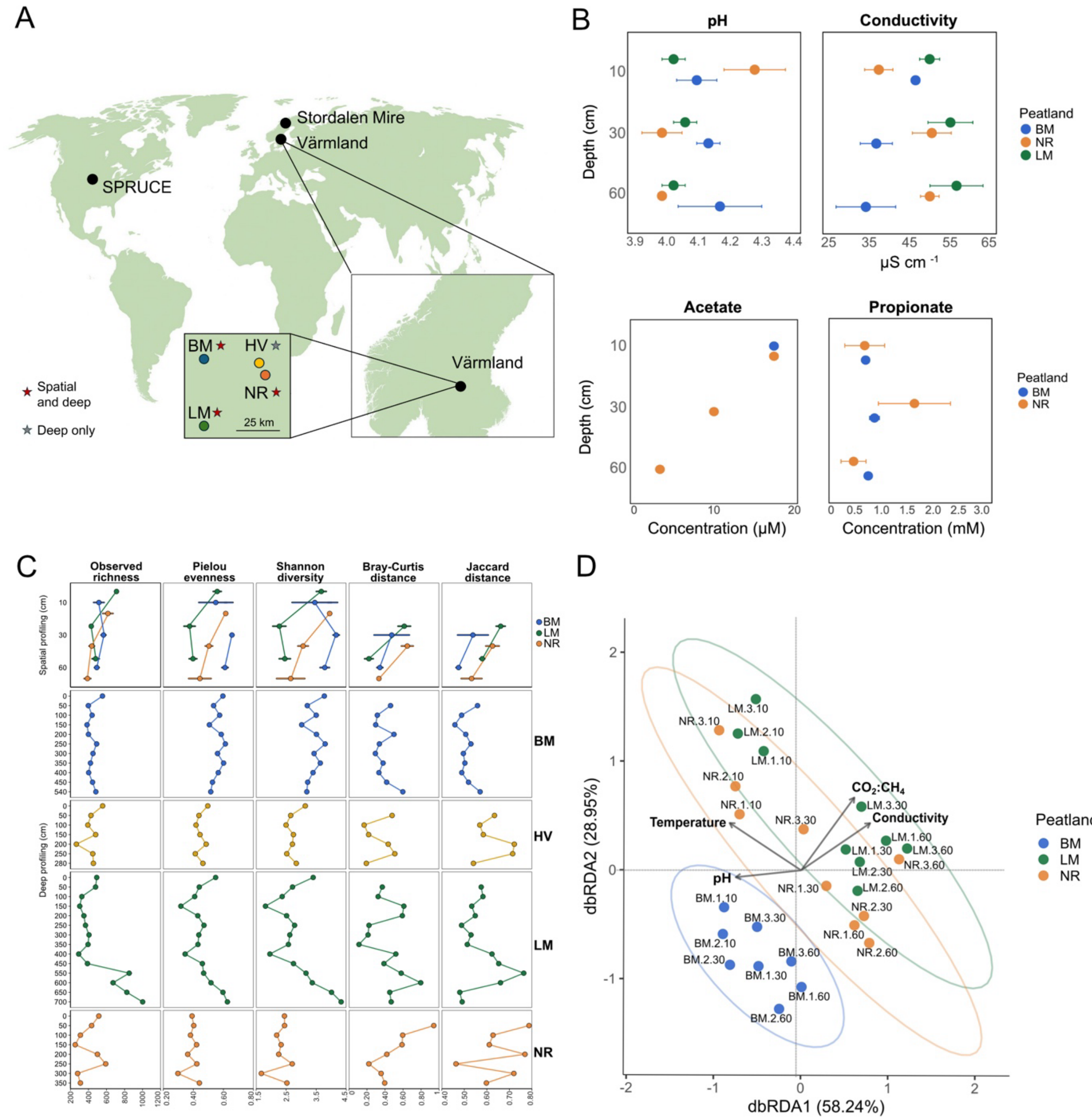


**ED Figure 1**. **Metadata and microbial community diversity. A**. Map showing the location of the Värmland sites (Björsmossen (BM), Norra Romyren (NR), Lungsmossen (LM), Havsjömossen (HV)) in relation to SPRUCE and Stordalen Mire. Red star "spatial and deep" indicates sites sampled for both spatial and deep profiling; grey star "deep only" highlights HV, sampled for deep profiling only. **B.** pH, conductivity (μS cm$^{-1}$) and organic acids (acetate (μM) and propionate (mM)) measured at 10, 30 and 60 cm depth shown as the mean across sites 1-3 for each peatland (BM: blue; NR: orange; LM: green). Horizontal bars indicate standard deviation values. **C**. 16S rRNA gene-based alpha diversity and beta diversity metrics (observed richness, Pielou evenness, Shannon diversity, Bray-Curtis and Jaccard distances) for spatial and deep profiling across peatlands (BM, HV, LM, NR). Points represent individual samples for deep profiling, while indicating mean values across sites (1-3) for spatial profiling. Bars indicate standard deviation. Bray-Curtis and Jaccard distances were calculated as dissimilarity to the immediately preceding depth within the same peatland, therefore beta-diversity values are absent for the first depth of each profile: 10 cm for spatial profiling and 0-25cm for deep profiling. **D**. Distance-based redundancy analysis (dbRDA) showing relationship between microbial community and biogeochemical measurements for BM, NR and LM in spatial profiling. Blue (BM), Green (LM) and orange (NR) points indicate peat microbial samples. while in the label above the points, 1,2,3 indicate sites and 10,30,60 the corresponding depth per sample. Arrows indicate environmental variables (pH, temperature, Conductivity, $CO_2$:$CH_4$).

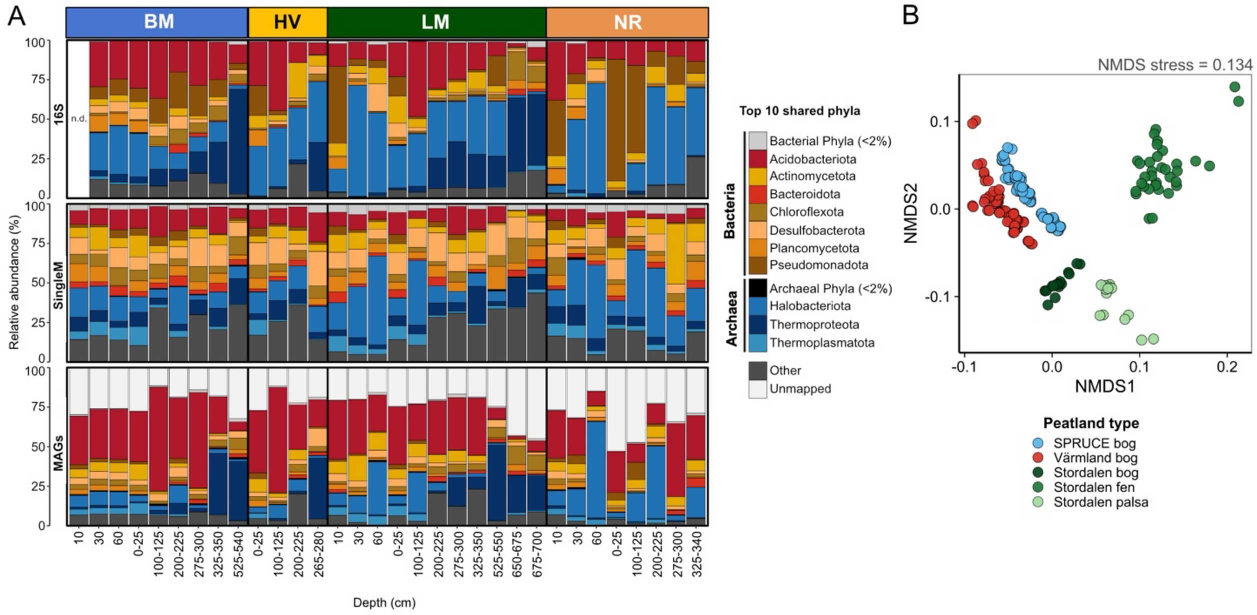


**ED Figure 2**. **Comparison of Värmland bog microbial communities with well-studied northern peatlands and across profiling approaches. A.** Comparison of microbial community composition across peatlands and depths using three approaches: 16S rRNA amplicon sequencing (top), taxonomic marker-gene-based profiling of raw metagenomic reads via SingleM (SingleM, middle) and metagenome-assembled genomes (MAGs) (MAGs, bottom). Stacked bar plots delineate the relative abundance (%) of the top 10 shared phyla across 32 metagenomic samples. Samples are grouped by peatland (BM, HV, LM, NR) and ordered by depth (cm). Phyla contributing <2% of relative abundance within each sample are grouped as *Bacterial Phyla (<2%*) or *Archaeal Phyla (<2%)*. Other phyla are grouped as *Other*, while metagenomic reads not mapping to MAGs are defined as *Unmapped*. n.d: no data. **B.** Non-metric Multidimensional Scaling (NMDS) ordination of MASH distances based on shotgun metagenomic reads comparing the Värmland bog sites (red) to SPRUCE bog (light blue) and Stordalen mire bog, fen, palsa (green). LM: Lungsmossen; NR: Norra Romyren; HV: Havsjömossen; BM: Björsmossen.

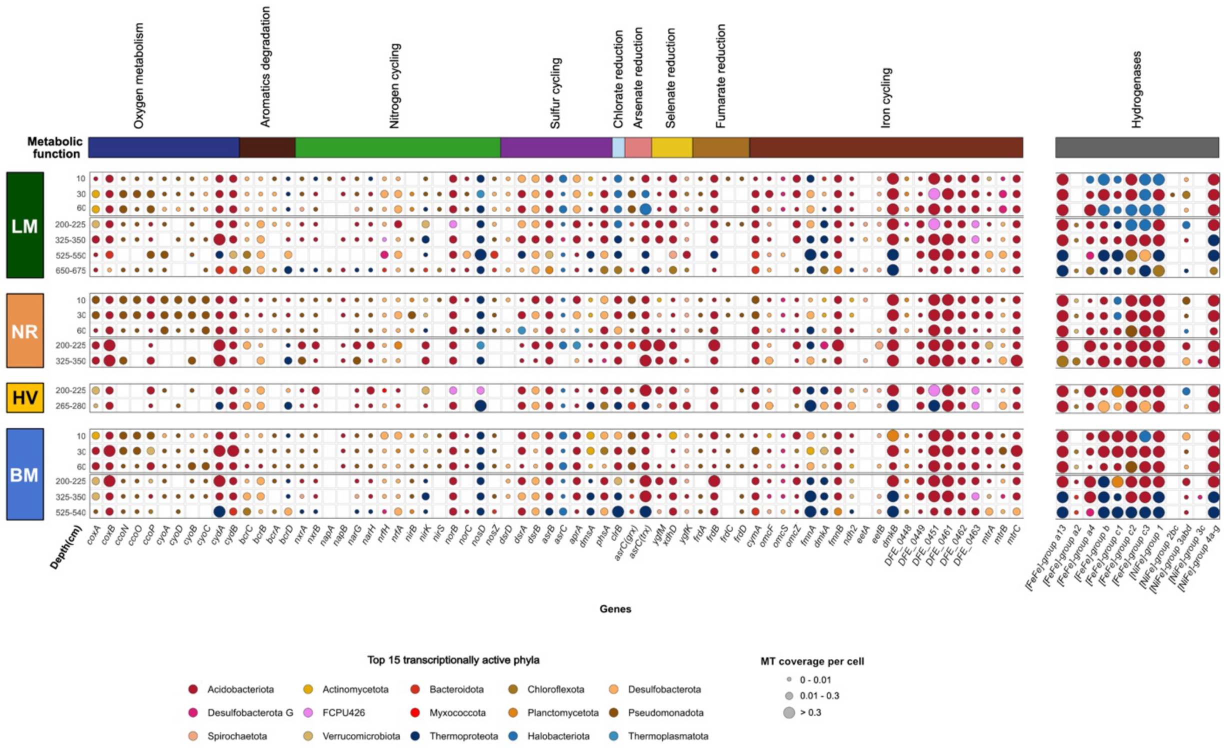


**ED Figure 3**. **Expression of microbial terminal reductases and hydrogenases across four Swedish sites and depths.** Bubble plot showing the expression of genes encoding microbial terminal reductases and hydrogenases across LM, NR, HV, BM and along the peat depth profile (cm). Individual terminal reductases are grouped by major metabolic functions indicated by the colored bar at the top (oxygen metabolism, aromatic degradation, nitrogen cycling, sulfur cycling, chlorate reduction, arsenate reduction, selenate reduction, fumarate reduction, iron cycling). Hydrogenase genes are grouped on the side and categorized according to defined HMMs. Bubble size reflects the level of expression classified into three categories (0-0.01, 0.01-0.3, >0.3), whereas bubble color indicates the most highly function-expressing phylum for each peatland x depth x function combination, within the 15 most transcriptionally active phyla globally. Absence of bubble indicates no detectable expression. LM: Lungsmossen; NR: Norra Romyren; HV: Havsjömossen; BM: Björsmossen.

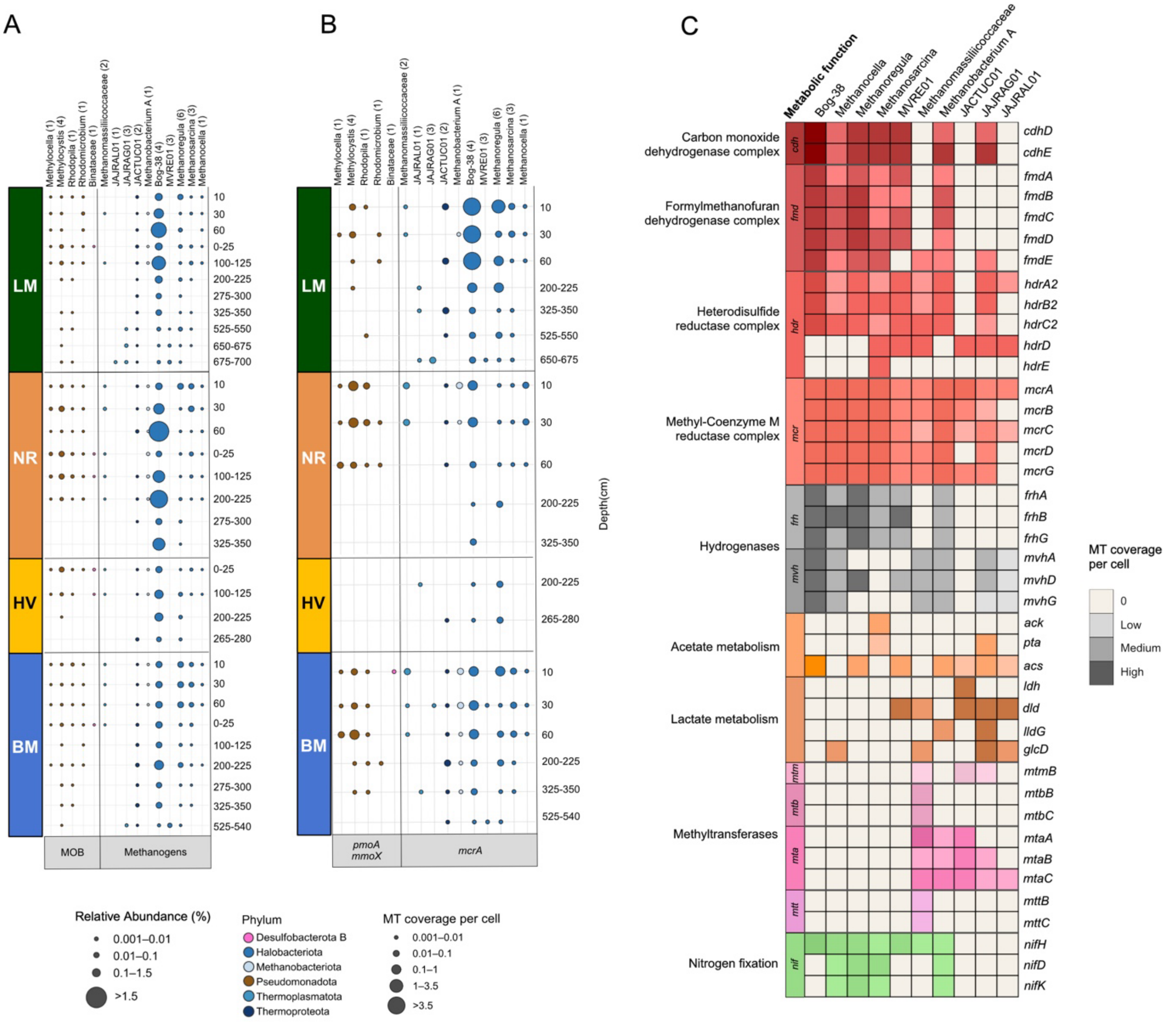


**ED Figure 4. The methane-cycling microbial diversity in the four Swedish bogs. A.** Identified methane-cycling MAGs and their relative abundances. The bubble plot shows the different microbial genera and families represented by the identified MAGs involved in methane cycling across the peatland sites (BM, HV, LM, NR) and depths (cm). Numbers in brackets next to the microbes' names indicate how many MAGs were assigned to each taxon. Circle colors denote the corresponding phylum, and circle sizes represent the relative abundance (%) of each MAG. Each bubble reflects the summed abundance of all MAGs belonging to a certain taxon. **B**. Expression levels of methane-cycling marker genes in the identified MAGs. Bubble colors follow the same phylum legend as in (**A**). Circle sizes represent the summed MT coverage per cell of each methane-cycling marker gene across all MAGs assigned to that taxon at each depth. **C**. Heatmap of pathway-specific marker genes expressed across all methanogens. MT coverage per cell values were classified into four categories (0, Low, Medium, High) with thresholds on the 25th and 75th percentiles of all non-zero expression values. Marker genes are grouped by metabolic functions and color gradients indicate the expression level of each gene. MOB: methane-oxidizing bacteria. LM: Lungsmossen; NR: Norra Romyren; HV: Havsjömossen; BM: Björsmossen.

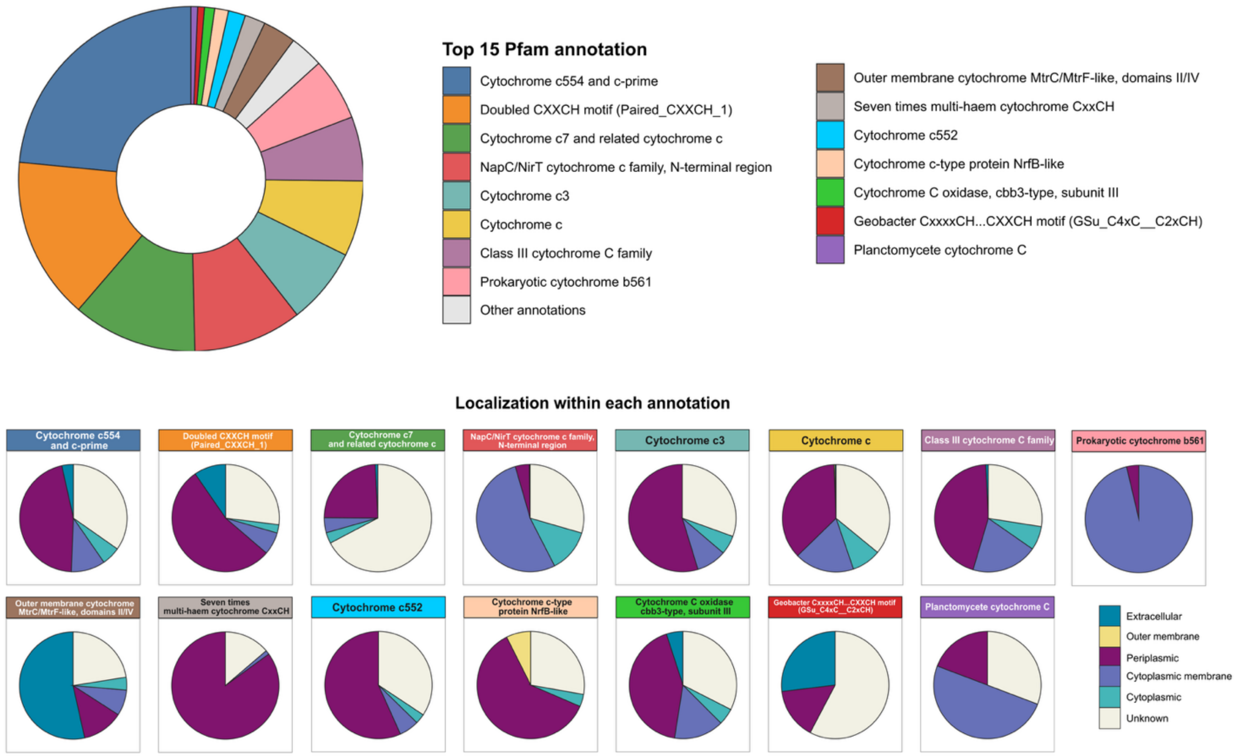


**ED Figure 5. Top 15 Pfam annotations among tier1 MHCs and predicted subcellular localization within each annotation class.** The donut chart summarized the top 15 Pfam annotations for tier1 (high-priority) MHCs and their predicted subcellular localization. Dominant annotations include cytochrome c544/c-prime (blue), doubled CXXCH motifs (orange) and cytochrome c7-related proteins (green). Sublocalization predictions reveal that most annotations are enriched in the periplasmic compartment, while specific families (e.g. NapC/NirT) are mainly associated with the cytoplasmic membrane and MtrC/MtrF-like cytochromes with the extracellular compartment, reflecting the functional specialization of MHCs across cellular compartments.

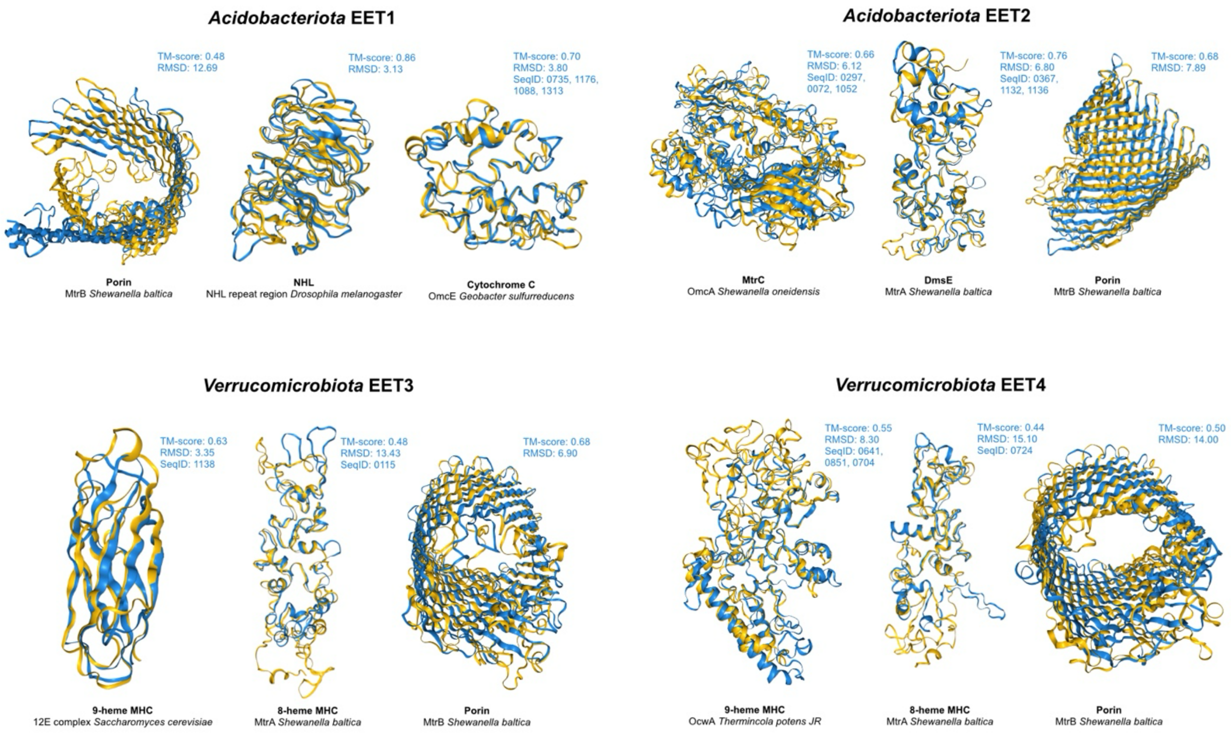


**ED Figure 6. Best structural hits of core EET proteins against PDB100 database (Foldseek).** The figure shows the structures of the core EET proteins (blue, query structures) and their best structural matches identified against the PDB100 database (yellow, target structures) via Foldseek search. Core protein names are shown in bold at the bottom, while corresponding matches and organisms are in regular font and italic. TM-score and RMSD values are reported for each match, together with the sequence clusters identifiers (SeqID) the core proteins belong to in our dataset.

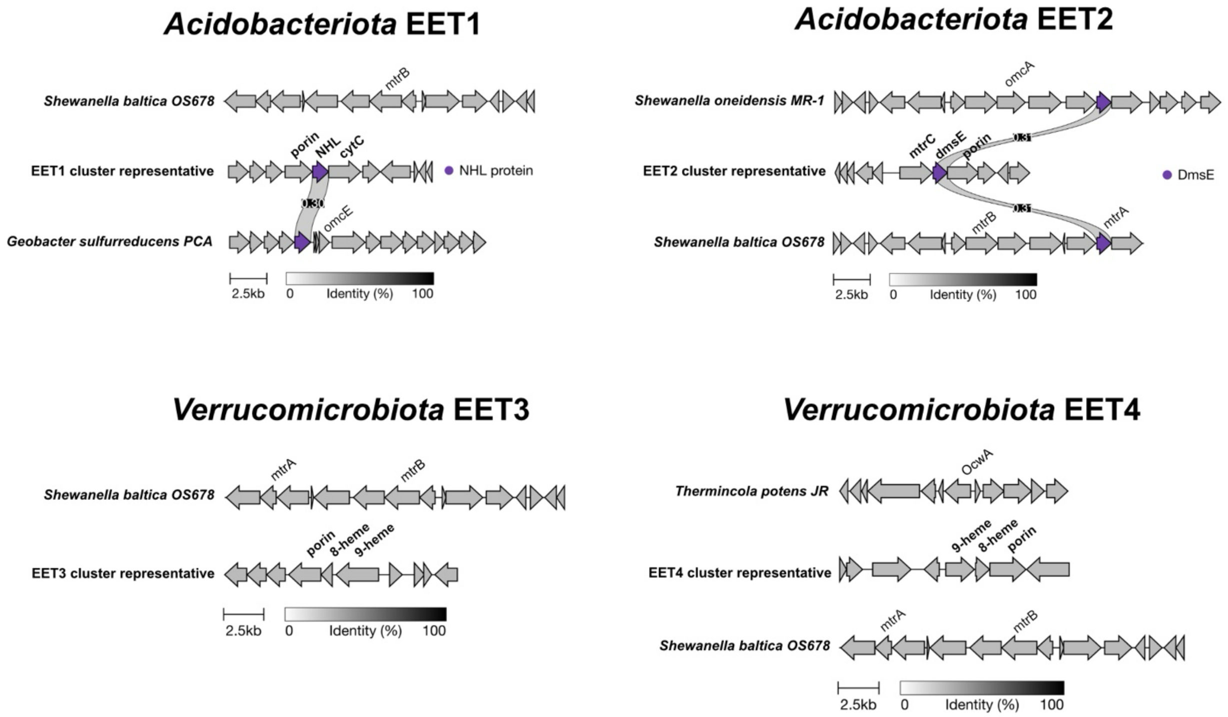


**ED Figure 7: conserved EET gene clusters share low sequence identity with model organisms.** Clinker alignments of representative EET clusters (EE1-EET4) core genes from Acidobacteriota and Verrucomicrobiota against genomic regions from reference organisms (*Shewanella oneidensis* MR-1*, Shewanella baltica* OS678*, Geobacter sulfurreducens* PCA*, Thermincola potens* JR), selected based on best structural matches of the encoded proteins. Sequence identity to reference genes remains low, close to the clinker identity threshold (-i 0.3), indicating strong sequence divergence.

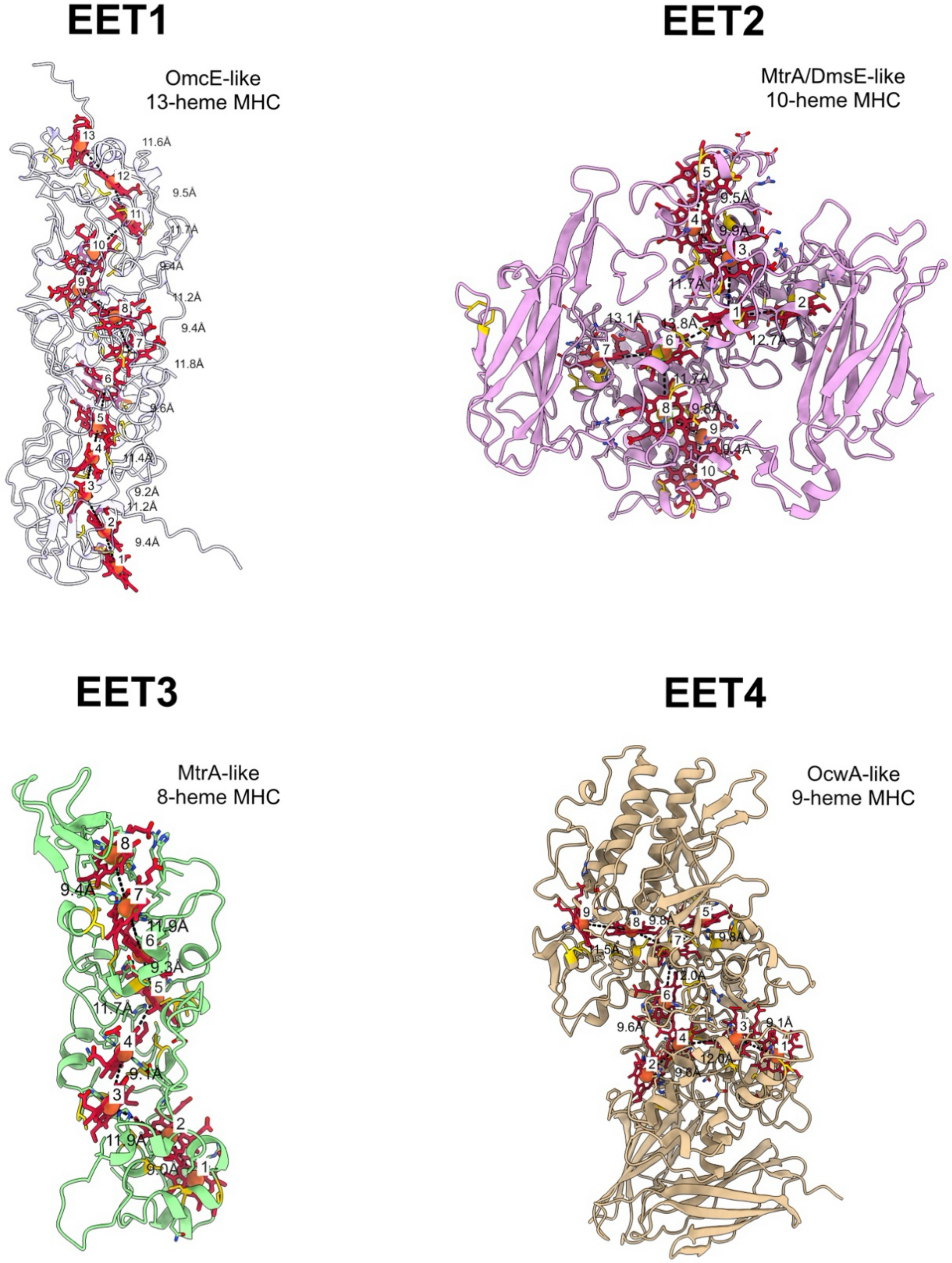


**ED Figure 8: multiheme cytochromes architecture supports electron hopping and potential extracellular electron transfer.** Structural models of core proteins from the representative EET clusters (EET1-EET4). Core MHC were modeled with heme ligands, with heme numbers inferred from CXXCH motifs. Redox centers are spaces within ~10-14 Å, likely enabling efficient electron hopping. These proteins are predicted to localize extracellularly (EET1, EET4) and in the periplasm (EET2, EET3).
