## Supplementary material for "Conserved multiheme cytochrome machinery for extracellular electron transfer is widespread and transcriptionally active across deep peat profiles": Fiorito_et_al_2026_Supplementary_information

This file contains:

**Supplementary Discussion**

1. Depth-dependent activity patterns across key microbial metabolisms
2. Uncultured Bog-38 dominates methanogenic activity while methanotrophic bacteria are active across depth
3. Details on colocalized genes in EET gene clusters
4. Complex organic matter degradation and $H_2$ cycling potentially fuel EET systems

**Supplementary Table**

Supplementary Table 6: overview of conserved EET clusters

**Supplementary Methods**

**Supplementary References**

### Supplementary Discussion

#### 1. Depth-dependent activity patterns across key microbial metabolisms

The expression of key metabolic pathways varied across the four Swedish peatlands and across depths.

Regarding **fermentation,** some pathways were highly expressed, especially those related to acetate metabolism, carrying the acetyl-coA synthetase (*acs*), acetate kinase (*ack*) and phosphate acetyltransferase (*pta*) genes (**Figure 2**). This likely reflected an active carbon turnover by microbial communities where acetate could be both produced and consumed, serving as energy and carbon source for other microbes[1]. Members of the Acidobacteriota phylum showed the highest expression of those genes across all peatlands and depths.

**Lactate metabolism**, also part of fermentation pathways, was predominantly expressed in the deeper peat layers (**Figure 2**). Genes encoding L-lactate dehydrogenase gene (*ldh*) were highly expressed, particularly by Thermoproteota in BM and LM (500-700 cm). Lactate has been reported to serve as energy and carbon source of cultivated methanogens belonging to the same phylum[2]. In addition, *ldh* expression was also observed in Verrucomicrobiota and Acidobacteriota, suggesting that these phyla may use lactate as electron donor.

**Nitrogen-cycling** genes (fixation, nitrite/nitrate/nitric oxide reduction, nitrification) were moderately expressed across depths, primarily by Acidobacteriota, Pseudomonadota and Halobacteriota. Nitrite reduction (nitrite reductase, *nrfA*) and nitrous oxide reduction (nitric oxide reductase, *norBC*) were expressed at depth by Desulfobacterota and Acidobacteriota, respectively (**ED Figure 3**).

Within the **C1 metabolism** macro-group, a higher expression of genes involved in aerobic carbon monoxide (CO) oxidation (*coxS, coxM, coxL*) was observed compared to other pathways (**Figure 2**). These genes were mainly associated with Thermoproteota and Pseudomonadota, suggesting that aerobic CO oxidation may represent a metabolically stable strategy also across peatlands, besides volcanic environments[3].

Genes involved in **oxygen metabolism** were also predominantly expressed across all peatlands and depths (**Figure 2**). Interestingly, Acidobacteriota expressed cytochrome *c* oxidases of both *bd*- and *caa*3-types. The *bd*-type oxidases function at sub-micromolar $O_2$ concentrations and contribute to the proton motif force through charge separation, i.e., consuming cytoplasmic protons for water formation while oxidizing quinol at the periplasmic side, without directly pumping protons[4]. *Caa*3-type oxidases are less $O_2$-sensitive than *bd*-type and couple the oxygen reduction (forming water) to vectorial ion transport, generating a proton motive force used for ATP production[5]. Although typically associated with aerobic respiration, expression of *bd*- and *caa*3-type oxidases in strictly anoxic layers likely reflects alternative roles, such as protection against toxic compounds (e.g., hydrogen peroxide, nitric oxide and hydrogen sulfide that may constitute other microbial metabolic intermediates) or oxygen scavenging, rather than aerobic growth[5]. Consistent with this, *bd*-type oxidases were upregulated under anoxic conditions, whereas *caa*3-types oxidases were associated with oxic conditions in Acidobacteriota enrichment cultures[6]. The high expression of caa3-type genes observed in anoxic layers in our dataset is unexpected but may reflect metabolic flexibility in Acidobacteriota*,* enabling them to switch their metabolism from anoxic to oxic conditions[6],5/26/26 9:42:00 PM potentially supported by transient oxygen availability generated by other microbial metabolisms. Such oxygen in deep peat layers may originate from cryptic production processes, including the so-called “dark oxygen” generation, a phenomenon recently hypothesized to occur in light-independent environments, such as the deep-seafloor and capable of generating oxic micro-niches[7].

**Hydrogenases** were actively expressed across all peatlands and depth, pointing to an active $H_2$ metabolism. Most of their expression was associated with Acidobacteriota, Thermoproteota and Halobacteriota (**Figure 2, ED Figure 3**), suggesting that these phyla play a central role in $H_2$ turnover in our studied peatlands. Halobacteriota expressed [FeFe]-hydrogenases belonging to the $H_2$-sensing groups c1-c3 (**ED Figure 3**). The precise function of these hydrogenases remains unclear, but they are thought to sense $H_2$ concentrations and regulate downstream pathways through associated regulatory elements, such as kinases and phosphatases[8]. Acidobacteriota showed the broadest expression of hydrogenase genes, with transcripts detected across all sites and depths (**ED Figure 3**). This included [FeFe]-hydrogenases of groups a1 and a3, involved in fermentation processes and reversible electron bifurcation from $H_2$ to ferredoxin and NAD, respectively[8], as well as regulatory [FeFe]-hydrogenases of $H_2$-sensing groups c1 and c3, energy-conserving [NiFe]-hydrogenases of groups 4a-g associated with $H_2$ production[9] and group 1 [NiFe]-hydrogenases involved in $H_2$ uptake. This combination suggests that Acidobacteriota may act simultaneously as $H_2$ producers and consumers, also utilizing $H_2$ as an electron donor. Consistent with this, members of this phylum have been shown to scavenge atmospheric $H_2$ under low-$O_2$ conditions for survival[10], suggesting $H_2$ may similarly support their persistence in low-$O_2$ deep peat layers. Thermoproteota dominated the expression of nearly all hydrogenase types described above, especially in the deepest layers (e.g LM 325-675 cm and BM 200-540 cm) (**ED Figure 3**). As this phylum also dominates community relative abundance at depth (according to SILVA taxonomy, Crenarchaeota, **Figure 1D**), the elevated hydrogenase expression may partly reflect this numerical dominance rather than enriched per-cell activity. Nevertheless, the breadth of expressed hydrogenase types suggests that $H_2$ metabolism plays a key role in sustaining Thermoproteota under likely energy-limited conditions in deep peat.

#### 2. Uncultured Bog-38 dominates methanogenic activity while methanotrophic bacteria are active across depth

Given the importance of methane cycling in northern peatlands, we examined the diversity, vertical distribution, and *in situ* expression of methane-cycling microorganisms across the peat profile.

No MAGs affiliated with anaerobic methanotrophic archaea (ANME), especially "*Candidatus* Methanoperedens" (ANME-2d), the lineage most clearly associated with freshwater ecosystems, were recovered. Therefore, we found no direct genomic evidence that archaeal anaerobic oxidation of methane is a notable natural methane mitigation process in these northern peatlands.

We detected several (predicted) methanogens including Halobacteriota *(Methanoregula, Methanocella, Methanosarcina,* MVRE01), Methanobacteriota, and less characterized Thermoplasmatota (e.g., JACTUC01) and Thermoproteota (e.g., JAJRAG01, JAJRAL01) genera (**ED Figure 4 A,B,C**). Collectively, these lineages encode the full spectrum of methanogenic pathways, i.e., hydrogenotrophic, methylotrophic (and potentially methoxydotrophic) and acetoclastic methanogenesis, fueled by $H_2$+$CO_2$, methylated compounds and acetate, as inferred from canonical marker genes (**ED Figure 4C**), consistent with multiomics studies from the further north Stordalen Mire peatland[11,12].

Among hydrogenotrophic methanogens, Bog-38*, Methanocella,* and *Methanobacterium A* expressed formylmethanofuran dehydrogenase (*fmd*) and heterodisulfide reductase (*hdr*) together with $F_{420}$ non-reducing hydrogenase (*mvh*) and the coenzyme $F_{420}$-reducing (*frh*) hydrogenases. Acetoclastic methanogenesis was mainly associated with *Methanosarcina*, which expressed carbon monoxide dehydrogenase/acetyl-CoA synthase subunits (*cdhDE*), acetate kinase (*ack)* and phosphotransacetylase (*pta*). Members of the Methanomassiliicoccaceae expressed a full suite of methyltransferases genes (*mtt, mta, mtb, mtm*), indicating that they actively utilized methylated compounds to fuel methanogenesis (**ED Figure 4C**).

Several methanogen MAGs also expressed traits indicative of broader metabolic flexibility (**ED Figure 4C**). Lactate, a common fermentative substrate also present in ombrotrophic peatlands[6], may provide an additional carbon and electron source for methanogenesis, via expression of genes encoding D-lactate dehydrogenase (*dld*), the L-lactate dehydrogenase complex protein (*lldG*), L-lactate dehydrogenase (*ldh*) and the glycolate dehydrogenase FAD-linked subunit (*glcD*) which happens to be the original annotation for the LDH that is part of an electron-confurcating complex[13]. Notably, Thermoplasmatota members (Methanomassiliicoccaceae*,* JAJRAG01, JAJRAL01) expressed both *glcD* and either *dld* or *lldG.* Lactate oxidation via *glcD* could provide electrons to *hdrD* for reduction of CoM-S-S-CoB[14]. Finally, several methanogens (especially *Methanocella, Methanoregula, Methanosarcina*) expressed nitrogen fixation genes (*nifHDK*), particularly at depth, providing a competitive advantage in nitrogen-limited environments[15].

#### 3. Details on colocalized genes in EET gene clusters

##### Overview and selection of cluster representatives

Four conserved EET-like gene architectures were identified across Acidobacteriota (EET1, EET2) and Verrucomicrobiota (EET3, EET4). To characterize the predicted functions of their constituent proteins, one representative MAG was selected per architecture for detailed manual annotation, comprising domain analysis, subcellular localization prediction, BLAST homology searches and AF3-based structural comparison.

Acido EET 1: NorraRomyren_4_0_25cm_2024_06_iMG_metabat2_bin.139

Acido EET 2: Lungsmossen_2_site3_30cm_2024_06_iMG_metabat2_bin.94

Verruco EET 3: Bjorsmossen_7_275-300cm_2024-06_iMG_metabat2_bin.72

Verruco EET 4: NorraRomyren_1_site3_10cm_2024_06_iMG_metabat2_bin.70

A summary of the core proteins, their predicted localizations, domain annotations and structural matches is provided in **Supplementary Table 6**. For each architecture, the conserved core proteins are described below.

##### Acidobacteriota clusters

To investigate conserved gene co-localization potentially supporting EET, we extracted the genomic loci of the extracellular MHC-encoding genes in Acidobacteriota MAGs (±5 genes up- and downstream of the MHC where possible), yielding 156 contigs originating from 111 MAGs. Clinker alignments of these genomic regions revealed that most regions clustered across two large clusters (dark red and green, **Figure 3**), with some outliers not clustering at all (9) and some smaller clusters comprising two or three genomic regions. Two highly conserved gene arrangements were identified: a three-gene core comprising a porin, an NHL-repeat domain protein and a cytochrome *c* protein, present in 49 MAGs, and a four-gene core comprising MtrC, DmsE, a porin and a cytochrome *c* protein in 28 MAGs.

##### Acidobacteriota EET1: the Porin-NHL-CytC cluster

We manually inspected the EET1 Acidobacteriota locus using the NorraRomyren_4_0_25cm_2024_06_iMG_metabat2_bin.139 MHC region as a representative of the EET1 cluster.

**MtrB-like porin.** The MtrB-like porin carries an N-terminal signal peptide (Sec/SPI) and is predicted to localize in the outer membrane (psort --negative: 6 extracellular, 3.6 outer membrane). BLAST search revealed that 91/100 hits fall within Terriglobia (best match: 67% amino acid identity), with four hits in Bacillati and three in candidate bacterial phyla. The AF3-predicted protein structure matches the outer membrane beta-barrel protein MtrB of *Shewanella baltica*, a core component of the transmembrane Mtr complex electron-conduit complex (PDB: 6R2Q; TM-score: 0.48, RMSD: 12.69 Å; Seq. Id: 8.5%, E-value: 8.61e-9; Prob.: 1) (**ED Figure 6**)[16]. The amino acid sequence identity of 8.5%, clearly indicates convergent evolution of structurally similar proteins with distinct sequences.

**NHL β-propeller scaffold protein.** The NHL β-propeller scaffold protein carries an N-terminal signal peptide (Sec/SPI) and is predicted to localize in the cytoplasmic membrane (pSORT --negative: 4.9 CM, 2.5 periplasmic, 2.5 extracellular). BLAST search revealed a 6-bladed beta-propeller domain (DUF5128; PF17170) with 91/100 top hits in Terriglobia (best match: 83% amino acid identity). SignalP detects a Sec/SPI signal peptide, consistent with export to the cell envelope/extracellular space. The AF3-predicted protein structure matches the NHL repeat region of *Drosophila melanogaster* (PDB: 6XG7; TM-Score: 0.86, RMSD: 3.13 Å, Seq. Id: 28%, E-value 1.99e-24; Prob.: 1.0) (**ED Figure 6**). The NHL-like domain (cd14962) is an uncharacterized NHL-repeat domain in bacterial proteins that resembles the WD repeat and other beta-propeller structures. This protein likely functions as a scaffold domain that may provide a binding interface for assembling a porin-cytochrome complex.

**13-heme OmcE-like MHC.** The 13-heme OmcE-like cytochrome *c* protein is predicted to be located extracellularly (pSORT --negative: 9.7 extracellular) and carries a signal peptide (Sec/SPI). BLAST search revealed homologues annotated as cytochrome *c*3 family proteins (PF14537), predominantly within Acidobacteriota (89/100 top hits; best match: 75% amino acid identity; 5 hits in Bacillati, 2 in candidate bacterial phyla, 1 in Verrucomicrobiota). The protein harbors 13 heme-binding motifs (7 canonical CXXCH, 3 $CXXC_nH$ and 3 $CX_nCH$). The AF3-predicted protein structure (ipTM = 0.92, pTM = 0.9) matches OmcE nanowires from *Geobacter sulfurreducens* (PDB: 7tfs, TM-score: 0.70, RMSD: 3.8 Å, Seq. Id: 29%, E-value 2.93e-8; Prob.: 1) (**ED Figure 6**). OmcE proteins form a filamentous appendage thinner than OmcS, but share a conserved heme-packing arrangement [17]; we similarly propose that this cytochrome *c* protein, as a core component of the Acidobacteriota EET1 cluster, could enable long-distance electron transport. Indeed, modeling the extracellular MHC of NorraRomyren_4_0_25cm_2024_06_iMG_metabat2_bin.139 together with 13 heme *c* ligands revealed that the Fe complexed within the heme *c* were consistently stacked at inter-iron distances of 9.2 to 11.8 Å, within the range permitting electron hopping between adjacent hemes.

**Acidobacteriota EET2: the MtrC-DmsE-Porin-CytC cluster**

The EET2 cluster representative is Lungsmossen_2_site3_30cm_2024_06_iMG_metabat2_bin.94, and comprises four conserved genes: *mtrC, dmsE* (double CXXCH), a *tonB*-type porin and *cytC*.

**10-heme MtrC-like MHC.** The 10-heme MtrC-like MHC carries an N-terminal signal peptide is predicted to be located extracellularly (pSORT --negative: 9.7 extracellular). BLAST search revealed 98/100 top hits within Terriglobia (90 in Bryobacterales; one hit at 100% amino acid identity), and identified a decaheme *c*-type cytochrome belonging to the OmcA/MtrC family (TIGR03507), which mediates Mn(III/IV) and Fe(III) reduction in *Shewanella*[18]. We detected 10 heme-binding motifs. The AF3-predicted protein structure (ipTM = 0.91, pTM = 0.91) matches OmcA of *Shewanella oneidensis* (PDB: 4LMH; TM-score: 0.66, RMSD: 6.12 Å, Seq. Id: 19%, E-value 7.65e-29, Prob.: 1.0) (**ED Figure 6**), the outer membrane decaheme cytochrome involved in EET[19].

**10-heme DmsE-like MHC** (double CXXCH). The 10-heme DmsE-like MHC carries a signal peptide and is predicted to localize in the periplasm (pSORT --negative: 9.8). BLAST search revealed 90/100 top hits within Terriglobia (80 in Bryobacterales; best match: 94% amino acid identity) and identified a decaheme *c*-type cytochrome belonging to the DmsE family domain (TIGR03508), which in *Shewanella* is implicated in extracellular respiration of dimethyl sulfoxide[20]. We detected 10 heme-binding motifs. The AF3-predicted protein structure (ipTM: 0.90, pTM: 0.83) matches MtrA from *Shewanella baltica* (PDB: 6r2a, chain A, TM-score: 0.76, RMSD: 6.8 Å, Seq. Id.: 31%, E-value: 1.07e-15; Prob.: 1.0) (**ED Figure 6**).

**MtrB-like porin (TonB-type).** The MtrB-like porin carries an N-terminal signal peptide and is predicted to localize in both the cytoplasmic and outer membrane (pSORT --negative: 4.7 cytoplasmic membrane, 4.8 outer membrane). BLAST search revealed 91/100 top hits within Terriglobia (78 Bryobacterales); no putative conserved domains or heme-binding motifs were detected. The AF3-predicted structure (ipTM: 0.83) matches the MtrB of *Shewanella baltica* (PDB: 6r2q, chain B; TM-score: 0.68, RMSD: 7.89 Å, Seq. Id: 11.9%, E-value: 1.49e-16; Prob.: 1.0) (**ED Figure 6**), which forms a 26-strand beta-barrel that shields the periplasmic 10-heme cytochrome MtrA in *Shewanella* from the membrane lipidic environment[16].

**4-heme periplasmic CytC**. A fourth cytochrome *c* is present in almost all EET2 clusters, adjacent to *mtrB* (**Figure 3C**). It carries a signal peptide, is predicted to localize in the periplasm (pSORT --negative: 9.8) and harbors 4 heme-binding motifs. The protein is conserved in Terriglobia (93/100 top BLAST hits, 83 of which are Bryobacterales) and possesses two TsdA domains (COG3258) encoding the C-terminal cytochrome *c* domain of thiosulfate dehydrogenase TsdA. The AF3 model (pTM: 0.72) yields no credible structural hit (best match: cytochrome *cd*1 nitrite reductase of *Pseudomonas aeruginosa*; Prob.: 0.41). We hypothesize that this 4-heme periplasmic MHC interacts with the Mtr-like complex, bridging the periplasmic space to complete the electron conduit.

#### Verrucomicrobiota clusters

Across the recovered 84 Verrucomicrobiota MAGs, 54 contigs encoded extracellular MHCs. Extracting and aligning the genomic loci bearing these MHCs with clinker revealed a conserved, syntenic gene neighborhood comprising several predicted *c*-type and *b*-type cytochromes, 9-heme CXXCH and 8-heme CXXCH motif proteins, and a porin (**Figure 3C**; annotated as MtrB/PioB-like (PF11854) and PEP-CTERM system-associated outer membrane protein (TIGR03016)).

##### *Verrucomicrobiota* EET3: the nonaheme MHC-octaheme MHC-MtrB-like porin cluster

The EET3 cluster is represented by Bjorsmossen_7_275-300cm_2024-06_iMG_metabat2_bin.72 (Palsa-1439, Verrucomicrobiota).

**Nonaheme MHC.** The first conserved gene encodes a nonaheme MHC with a signal peptide that is predicted to localize extracellularly (pSORT --negative: 9.7). BLAST search revealed 53/100 top hits within Verrucomicrobiota, 14 in the FCPU426 phylum, 13 in Acidobacteriota, 7 Chloroflexota and a few others. Three functional domains were detected: an UshA-like domain (NF041198), a myxosortase domain (NF038112) and a tetraheme cytochrome *c* subunit NapC-type domain associated with nitrate or TMAO reductases (COG3005). The presence of a myxosortase-dependent M36 family metallopeptidase domain (NF038112) is notable, as this domain has been associated with outer membrane vesicle formation in some Verrucomicrobiota[21] and the protein is also found in Terriglobia (particulary Bryobacteraceae) Acidobacteriota, suggesting a potential link between outer membrane vesicle or appendage formation and EET that could extend the effective reach of electron transfer in soil systems. The AF3-predicted protein structure of the nonaheme MHC (pTM: 0.8, ipTM: not determined) best matches the Asf1-Fab complex of *Saccharomyces cerevisiae* (PDB: 6az2, chain D; TM-Score 0.63, RMSD: 3.35 Å, Seq. Id: 11.8%, E-value: 8.71e-4, Prob.: 1.0) (**ED Figure 6**), which serves as a histone chaperone[22]. Yet the low score likely makes any further interpretation on the structure or structural homology difficult.

**Octaheme MHC (double CXXCH).** The second conserved gene encodes an octaheme MHC with a signal peptide that is predicted to localize in the periplasm (pSORT --negative: 9.5). BLAST search revealed 68/100 top hits within Verrucomicrobiota, 10 in Thermodesulfobacteriota, 7 Acidobacteriota, 5 in Bacillati and some in others. We counted 8 heme-binding motifs (7 canonical, 1 non-canonical) and identified a DmsE family domain (TIGR03508) which in *Shewanella* is implicated in extracellular respiration of dimethyl sulfoxide[20]. The AF3-predicted protein structure (pTM: 0.82) matches MtrA from *Shewanella baltica* (PDB: 6r2q, chain A; TM-score: 0.48, RMSD 13.43 Å, Seq. Id: 26.4%, E-value: 1.51e-10; Prob. 1.0) (**ED Figure 6**).

**MtrB-like porin.** The third conserved gene encodes an MtrB-like porin with a signal peptide that is predicted to localize in the outer membrane (pSORT --negative: 9.5). BLAST search revealed 68/100 top hits within Verrucomicrobiota, 16 in Thermodesulfobacteriota, 6 in Myxococcota and a few others. A GSU2205 family CXXCH-containing domain (NF0410207) was detected; yet manual inspection did not reveal a heme binding motif (nor a premature stop codon indicative of recoding to selenocysteine, as sometimes associated with this domain), suggesting shared architectural features but a distinct function for this domain in the present protein. The AF3-predicted protein structure (ipTM : 0.89) matches MtrB from *Shewanella baltica* (PDB: 6r2q, chain B; TM-score: 0.68, RMSD: 6.9 Å, Seq. Id: 12.7%, E-value: 3.64e-15, Prob.: 1.0), which forms a 26-strand beta-barrel that accommodates the 10-heme cytochrome MtrA in *Shewanella* from the membrane lipidic environment[16] (**ED Figure 6**).

**Verrucomicrobiota EET4: the OcwA-like MHC-octaheme MHC-PEP-CTERM porin cluster**

The EET4 cluster is represented by NorraRomyren_1_site3_10cm_2024_06_iMG_metabat2_bin.70 (Limisphaerales, Verrucomicrobiota).

**Nonaheme OcwA-like MHC.** The first conserved gene encodes a 583 amino acid nonaheme OwcA-like MHC (cytochrome *c*-552/4 family) with a signal peptide that is predicted to be in the cytoplasmic membrane or extracellularly (pSORT --negative: 4.7 cytoplasmic membrane, 4.6 extracellular). BLAST search revealed 78/100 top hits within Verrucomicrobiota, 3 in Terriglobia, 7 in Bacillati and some in other bacterial taxa. It also identified an UshA-like protein domain (NF041198), which is distantly related to hydrolases. We detected 9 canonical CXXCH heme-binding motifs. The AF3-predicted (ipTM: 0.97, pTM: 0.95) matches the outer cell wall cytochrome OcwA from the thermophilic Gram-positive bacterium *Thermincola potens* JR (PDB: 6i5b, chain B; TM-Score: 0.55, RMSD: 8.3 Å, Seq. Id: 17.7%, E-Value 2.00e-10, Prob.: 1.0) (**ED Figure 6**). OcwA is a multipurpose terminal reductase capable of reducing solid electron acceptors, soluble electron shuttles and oxyanions[23].

**Octaheme MtrA-like MHC (double CXXCH).** The second conserved gene encodes a 286 amino acid octaheme MtrA-like MHC with a signal peptide that is predicted to be located in the periplasm (pSORT --negative: 9.8). BLAST search revealed 78/100 top hits within Verrucomicrobiota and identified a DmsE family domain (TIGR03508), also found in Acidobacteriota EET2 and Verrucomicrobiota EET3. We detected a total of 8 heme binding motifs (7 canonical and 1 $CX_4CH$). The AF3-predicted structure (ipTM: 0.93, pTM: 0.9) matches the MtrA from *Shewanella baltica* (PDB: 6r2q, chain A; TM-score: 0.44, RMSD: 15.1 Å, Seq. Id.: 25.5%, E-value 3.73e-9, Prob.: 1.0) (**ED Figure 6**).

**PEP-CTERM-associated outer membrane porin.** The third conserved gene encodes a 736 amino acid outer membrane porin (TIGR03016; PEP-CTERM system-associated outer membrane protein family) that is predicted to be located in the outer membrane (pSORT --negative: 9.5). BLAST search revealed 84/100 top hits in Verrucomicrobiota and identified a CirA domain (COG1629), which is an outer membrane TonB-dependent receptor that may facilitate import of extracellular nutrients, including Fe. The AF3-predicted protein structure (pTM: 0.86, ipTM: not determined) matches MtrB from *Shewanella baltica* (PDB: 6r2q, chain B; TM-score: 0.5, RMSD: 14.0 Å, Seq. Id: 11.2%, E-value 1.45e-14, Prob.: 1.0) (**ED Figure 6**).

#### 4. Complex organic matter degradation and $H_2$ cycling potentially fuel EET systems

Since conserved EET genes were expressed in both spatial and deep profiles (**Figure 4C**), we reconstructed the metabolism of Acidobacteriota (96 MAGs) and Verrucomicrobiota (24 MAGs) carrying the EET clusters (see **Methods**) to understand which energy and carbon sources may support their metabolism and where the electrons feeding these systems may come from (**Figure 5A,B**). This question was relevant especially in deep peat, where EET genes remained expressed despite the more redox-stable and oxygen-limited conditions expected at depth (**Figure 4C**).

Overall, EET-bearing microbes appeared to be mainly heterotrophic, with expressed Carbohydrate-Active Enzymes (CAZymes) indicating that polysaccharide degradation feeds sugars into glycolysis and the Entner-Doudoroff pathway, generating pyruvate. In addition, some MAGs carrying the EET systems fully expressed genes belonging to the THF-methyl branch of the Wood-Ljungdahl pathway (WLP), while microbes carrying EET1, EET2 and EET4 also encoded the carbonyl branch, suggesting that they could potentially be acetogens and grow autotrophically on $CO_2$. This module is linked to C1 metabolism via encoded and expressed methyltransferases for methylotrophy suggesting that methylated compounds may provide additional energy and carbon inputs. Fermentation products may also contribute to this network, as lactate-metabolism genes were expressed in EET1 and EET3 (*ldh*: L-lactate dehydrogenase) and EET1, EET2, EET4 (*ldhA*: D-lactate dehydrogenase) microbes. This occurred along with butyrate metabolism genes for EET1 and EET2 microbes (*buk*: butyrate kinase; *ptb*: phosphate butyryltransferase), while genes linked to acetate (*ackA*: acetate kinase; *acs*: acetyl-CoA synthetase; *pta*: phosphate acetyltransferase for EET2 and EET4 microbes only) and alcohol metabolisms (*ALDH*: aldehyde dehydrogenase, *adh*: alcohol dehydrogenase) were found across all EET-bearing MAGs. In addition, EET1 MAGs uniquely showed the potential for anaerobic aromatic degradation, suggesting that more complex peat-derived compounds may also serve as carbon sources.

Additional metabolic functions were associated with specific EET systems. Acidobacteriota carrying EET1 and EET2 encoded and expressed genes involved in dissimilatory sulfate reduction (*sat*: sulfate adenylyltransferase; *aprAB*: adenylylsulfate reductase subunits A and B; *dsrAB*: dissimilatory sulfite reductase alpha and beta subunits), together with the expression (EET1 only) of *betC* (choline sulfatase, **Figure 2**) indicating involvement in the sulfur cycling[24].

The two phyla also differed in their cytochrome C maturation machinery. Acidobacteriota expressed a broader set of maturation genes associated with the cytochrome c biogenesis machinery described in *Shewanella oneidensis*[25] including many cytochrome c-type biogenesis genes (*ccmA,B,C,E,F,G,H)*, the periplasmic monoheme cytochrome c5 (*ScyA*) and the Cytochrome c-type biogenesis thioredoxin (*SO0269*), whereas Verrucomicrobiota encoded a reduced subset (*ccmA,G scyA, SO0269*). This suggests that both lineages likely have the capacity to assemble and maintain multiheme cytochromes, although Verrucomicrobiota may rely on a simpler maturation system or on still uncharacterized components.

Together, these results support a model in which EET-bearing Acidobacteriota and Verrucomicrobiota sustain a heterotrophic lifestyle, mainly using organic matter and hydrogen as carbon and electron sources respectively, respiring aerobically when $O_2$ is available and redirecting electrons towards extracellular TEAs (potentially POM) through EET systems when $O_2$ or other soluble electron acceptors become limiting.

### Supplementary tables

**Supplementary Table 6. Overview of conserved EET clusters.**

Protein names are mostly derived from BAKTA annotations in genbank file. The name is the curated name based on structural homology and/or domain similarity. The localization is predicted with pSORT(--negative). The SeqC clusters show the cluster name of the member and other homologues from the clinker alignment. The number of Cluster members is shown in parentheses. The taxa that the EET cassettes were found are indicated on the order level and the number of MAGs recovered are found in parenthesis.

| Cassette | Protein | Name | Length (aa) | Localization | Pfam domains | SeqC | Structural match | Taxa |
|---|---|---|---|---|---|---|---|---|
| **A-EET1** | Porin | Porin | 613 | Outer membrane | | - | MtrB (PDB 6R2Q_B) | Terriglobales, UBA7540 |
| | NHL | NHL β propeller | 366 | Extracellular | NHL, 2 DUF5128 | - | (PDB 6XG7) | Terriglobales, UBA7540 |
| | Cytochrome c domain protein | 13-heme OmcE-like (CytC) | 726 | Extracellular | 2 Paired | 0735(51), 1176 (3), 1088 (9), 1313 (6) | OmcE (PDB 7tfs) | Terriglobales, UBA7540 |
| **A-EET2** | OmcA/MtrC | 10-heme OmcA-like (MtrC) | 740 | Extracellular | 2 Mtrc-MtrF_II-IV_dom | 0297(48), 0072 (2), 1052 (3) | OmcA (PDB 4LMH) | Bryobacterales |
| | Doubled CXXCH | 10-heme MtrA-like (DmsE) | 325 | Periplasm | 3 Paired_CXXCH | 0367(23), 1132 (13), 1136 (6) | MtrA (PDB 6r2a) | Bryobacterales |
| | MtrB/PioB | Porin | 708 | Cytoplasmic membrane/outer membrane | | - | MtrB (PDB 6r2q_B) | Bryobacterales |
| **V-EET3** | 9-heme MHC | 9-heme MHC | 883 | Extracellular | | 1138(6) | PDB 6az2 | Palsa-1439 |
| | Doubled CXXCH | 8-heme MtrA-like | 230 | Periplasm | 2 Paired_CXXCH | 0115(19) | MtrA (PDB 6r2a_A) | Palsa-1439 |
| | Porin | | 668 | Outer membrane | | - | MtrB (PDB 6r2q_B) | Palsa-1439 |
| **V-EET4** | 9-heme | 9-heme OwcA-like MHC | 583 | Extracellular/cytoplasmic membrane | | 0704(8), 0641 (4), 0851 (17) | OwcA (PDB 6I5B_B) | Limisphaerales |
| | Doubled CXXCH | 8-heme MtrA-like MHC | 286 | Periplasm | 2 paired_CXXCH | 0724(25) | MtrA (PDB 6r2q_A) | Limisphaerales |
| | TIGR03016 family PEP-CTERM systems-associated outer membrane protein | Porin | 736 | Outer membrane | | - | MtrB (PDB 6r2q_B) | Limisphaerales |

#### Supplementary Methods

##### ***Field sites and sampling***

These sites were previously characterized in long-term biogeochemical studies[26–30]; the four bogs are hydrologically isolated, receiving water exclusively from precipitation. Their vegetation is dominated by *Sphagnum* carpets and hummocks with few isolated trees (mostly *Pinus sylvestris*), consistent with earlier descriptions of ombrotrophic peatland sites in the area[31].
